# Exploring the potential for scanning electron microscopy/focused ion beam-based diffraction for screening cryo-transmission electron microscopy samples

**DOI:** 10.1101/2025.09.25.678486

**Authors:** Eric V. Woods, Christoph Wigge, Yujun Zhao, René de Kloe, Tim M. Schwarz, Ben Britton, Stefan Zaefferer, Baptiste Gault

**Affiliations:** Max Planck Institute for Sustainable Materials, Max-Planck-Str. 1, 40239 Düsseldorf, Germany; Electron Imaging Solutions EIS GmbH, Heinrich-Heine-Garten 10, 40549 Düsseldorf, Germany; EDAX / Gatan, Ringbaan Noord 103, 5046AA Tilburg, The Netherlands; Department of Materials Engineering, University of British Columbia, 309-6350 Stores Road, Vancouver, BC V6T 1Z4, Canada; University Rouen Normandie, INSA Rouen Normandie, CNRS, Normandie University, GPM UMR 6634, F-76000 Rouen, France

## Abstract

The study of biological and organic materials at high resolution using cryogenic transmission-electron microscopy (cryo-TEM) necessitates vitrification to preserve the native structure. Assessing sample integrity is essential, particularly as ice crystallization during freezing and handling can cause irrecoverable structural damage. Usually, a secondary cryo-TEM is used for initial screening, only possible after a time-consuming sample preparation workflow. In the present work, we propose simple methods that exploit existing workflows developed for materials science analyses and demonstrate on-grid *in situ* assessment of ice crystallinity with electron backscatter diffraction (EBSD) on a direct electron detector (DED) in a cryo-scanning-electron microscope (SEM). This evaluation step can be performed prior to sample preparation for cryo-TEM by using cryogenic focused ion beam (cryo-FIB) milling. Custom grid holders and jigs were developed to integrate the clipped cryo-TEM grids and evolve the sample preparation workflow. EBSD detects hexagonal ice in some areas of the samples, whereas other areas show an absence of EBSD signal, consistent with vitreous ice, that enable targeting the further steps of sample preparation for cryo-TEM. Off-axis transmission Kikuchi diffraction (TKD) was attempted, but led to severe damage to polished TEM-lamellae and appears unsuitable. A proof-of-concept lift-out from a clipped cryo-TEM grid mounted on a support is introduced, demonstrating possibilities for expanded cryogenic correlative workflows beyond the acceleration of sample screening for cryo-TEM.

## Introduction

The development of cryogenic transmission electron microscopy (cryo-TEM) in parallel with water vitrification techniques has revolutionized the structural analysis of biomolecules, and has enabled the determination of the structures e.g. of proteins and virus envelope in their native hydrated state ^1–5^ without requiring molecular crystallization and subsequent X-ray diffraction (XRD)^1,6,7^. Sample preparation protocols have evolved, but fundamentally to achieve water vitrification, i.e. creating amorphous water ice, cooling rates in excess of 10^5^ K/s are required to avoid damage from the volume expansion arising from the formation of crystalline cubic I*_c_* or hexagonal I*_h_* ice ^8,9^. Vitrification is most commonly achieved by plunge freezing TEM grids containing a solution with the biomolecules for single-particle imaging into fast-cooling liquid cryogens, e.g. ethane and propane^10^, as the Leidenfrost effect prevents achieving the necessary cooling rates with liquid nitrogen (LN2). For yeast, small organisms, etc. alternative methods are employed such as metal-mirror (“slam”) or high-pressure freezing (HPF) ^11–13^.

After freezing, a Cu ring is clamped around the edge of the TEM grid for more robust grid handling and to allowed automated transport ^14^. This procedure is referred to as “clipping”. However, clipping and the subsequent part of vendor-specific workflows are not readily compatible with existing holders and equipment designed for thin standard TEM grids that are unable to handle these thicker grids ^15^, requiring dedicated equipment.

For more complex samples, e.g. cells, the sample need to first be sectioned into electron-transparent regions of interest (ROIs). Historically, frozen samples would be cut into ultrathin slices with a diamond knife using a cryogenic ultramicrotome and individual slices transferred onto TEM grids ^8,12,16^. More recent methods can involve transfer of the grid or frozen specimen under cryogenic conditions to a Ga^+^ or plasma focused-ion beam (e.g. Xe^+^) (FIB / PFIB) dual-beam system equipped with a scanning electron microscope (SEM) and cryogenic sample stage ^17–19^. In this workflow, points of interest are localized using SEM imaging, correlation with e.g. light, ion and electron microscopy (CLIEM) ^20^, and combinations of different techniques ^15^. Traditionally, the FIB cuts and thins an electron-transparent lamella in particular ROIs on a TEM grid before transfer to cryo-TEM^17^. For samples prepared by HPF, site-specific preparation can be performed for a targeted region, for instance within a cell. A thicker slice is first cut, attached to a cryogenically cooled micromanipulator, lifted out, and transferred to a separate TEM grid, prior to final thinning to electron transparency (“cryo lift-out”) ^19,21,22^. This is particularly useful for samples prepared for TEM tomography in which a tilt series is taken to enable three-dimensional (3D) volumetric reconstruction ^23,24^. These workflows can be partially or fully automated in many cases ^19,25^.

After vitrification, robust and careful specimen handling and transfer are essential to avoid the devitrification of amorphous ice in case the temperature rises above -135°C^26^. The formation of crystalline ice can damage the samples to be investigated. Conventionally, samples are transferred into a lower-voltage cryo-TEM for initial screening to determine if the freezing process was successful, e.g. if the ice is crystalline or vitreous, before transfer to a cryo-FIB for further preparation or a cryo-TEM with higher resolution for data acquisition. Screening cryo-TEM grids consumes valuable microscope time; although crystalline samples are typically discarded because the specimens suffer freezing damage, seemingly intact regions may still be damaged, which would normally only be recognized after full lamella fabrication or imaging in the high-resolution imaging TEM.

In principle, ice crystallinity could be assessed in the SEM/FIB with *in situ* electron diffraction (ED), where the electron beam enters the specimen and diffraction patterns are collected of either the resultant backscattered or transmitted electrons. In electron backscatter diffraction (EBSD), electrons enter the sample and, following potentially multiple scattering events, some exit from the sample surface to form a diffraction pattern on an EBSD camera or detector placed a few degrees off axis, e.g. 90°, from the electron optical axis, with the sample highly tilted at an angle of 70° with the incident electron beam ^27^. For thin samples, in the range of a few hundreds of nanometers, it is also possible to collect electrons that have been transmitted through the sample, which can also form a diffraction pattern; this technique is called transmission Kikuchi diffraction (TKD)^28^. For TKD, the detector can be in the same position as for EBSD, in an ‘off-axis’ configuration, but may be under the sample for ‘on-axis’ TKD, close to a scanning transmission-electron microscope (STEM) configuration^29^. EBSD on water ice was previously demonstrated ^30–34^. High electron doses can however be required for ED, which can significantly damage even actively cryogenically cooled specimens^29,35^, as also observed in the TEM^26,36^.

New generation EBSD / TKD detectors using direct electron detectors (DEDs) have dramatically increased detector sensitivity, i.e. the collection efficiency of detected electrons as well as the angular acuity of the diffraction patterns, which allow the use of substantially reduced SEM accelerating voltages and total electron doses ^37–41^. The advent of commercially available DED-based EBSD systems merits reevaluation of the feasibility of using EBSD and/or TKD for rapid assessment of the crystallinity of vitrified samples on TEM grids prior to and during specimen fabrication. However, implementation of these new workflows faces practical challenges because of specific sample orientations, tilt angles between sample and detector etc., which cannot be reached without the development of new and dedicated sample holders^29^.

Herein, we demonstrate a simplified, unified workflow compatible with standard cryo-electron microscopy laboratory workflows, e.g. vitrified, clipped grids can be taken from a liquid nitrogen (LN_2_) storage dewar, transferred into the cryo-FIB through a low-moisture N_2_-filled glovebox and ultrahigh-vacuum (UHV) suitcase transfers^42–45^. To streamline the process, custom holders were designed that can carry TEM full grids or half-grids. The crystalline or amorphous nature of the ice can be assessed using EBSD, thereafter standard cryo-EM FIB sample preparation protocols for cryo-TEM are used. Besides EBSD, off-axis TKD of TEM grids prepared by plunge freezing is demonstrated, although it proved unsuitable for the assessment of the crystallinity of lamellae due to damage. Thereafter, the prepared grids are removed and subsequently transferred back to storage and later assessed and imaged using cryo-TEM. Our work demonstrate that the use of a fully UHV-cryogenic workflow allows for clean preparation and transfer of grids and on-grid lamellae, and that ED inside the SEM/FIB can be used to screen lamellae for future use in cryo-TEM.

### Experimental workflow

The overall workflow for sample preparation, fabrication, and validation is summarized in **Figure 1**.

**Figure 1.**
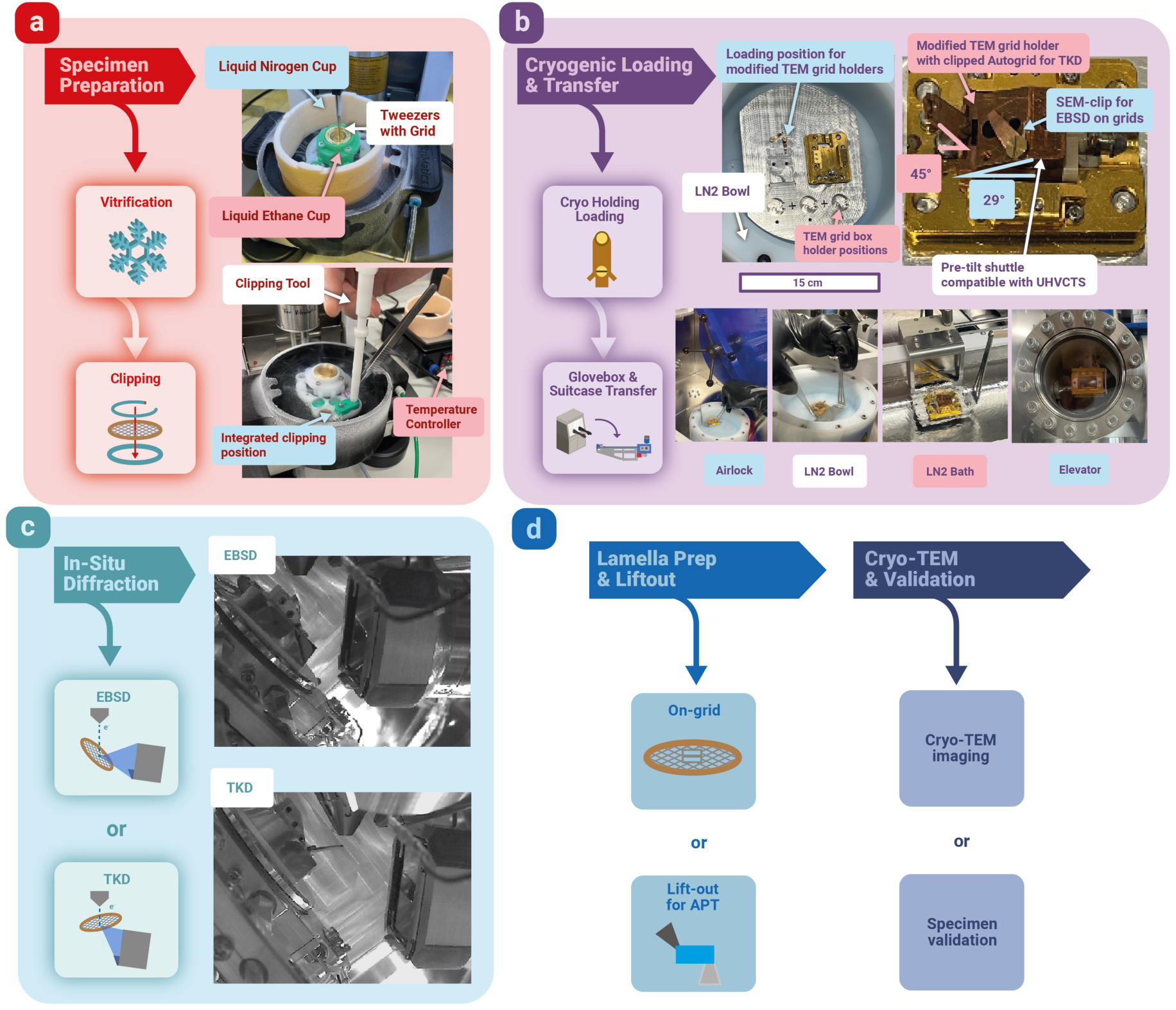
General workflow: (a) a Vitrobot with a VitriClip was used for vitrification of sea urchin sperm cells. Enlarged view of the bowl with integrated ethane condenser, clipping tool and temperature controller; (b) custom metal loading plate, with positions for custom TEM grid holder loading, grid boxes and the shuttle. Enlarged view of the loading plate and a mounted shuttle with the custom grid holder and the sample clip. Typical loading steps are shown with loading through the airlock into the glovebox, transferring the loading plate into the LN2 bath, and loading into the elevator for the UHV cryo transfer; (c) for EBSD analysis a cryo TEM grid is mounted with a 29° pre-tilt on the shuttle using a clip mechanism and tilted to 41° (70° total tilt) using the SEM stage. For TKD the TEM grid is mounted in the custom grid holder with a 45° pre-tilt on the shuttle; (d) on the TEM grid lamellas can be prepared as well as site specific lift out for APT. After clipping with the integrated clipping tool, the grids were placed into grid boxes in LN2 storage dewars until usage.

### Sample preparation

Sea urchin sperm cells in artificial seawater (10mM 2-(N-morpholino) ethanesulfonic acid, 200mM NaCl, 10mM CaCl_2_ with a pH of 6.5) was selected as a model sample. 3 µL of this solution were deposited on 1.2/1.3 µm Cu grids from Quantifoil (Quantifoil Micro Tools, Jena, Germany) that had been glow discharged for 10 s prior to use. The grids were vitrified using a Vitrobot Mark 4 (Thermo Fisher Scientific – TFS, Eindhoven, Netherlands) equipped with a VitriClip (Vitrimatics Maastricht, The Netherlands), shown in **Figure 1(a)**. Vitrification was performed at 4 °C and 100% relative humidity using a blot force of 0 and a blot time of 5 s. After clipping with the clipping tool, the grids were placed into grid boxes in LN2 storage dewars. Grids were retrieved from a long term LN2 storage dewar some weeks later.

### Cryogenic Loading and Transfer

#### Sample holders and sample transfer

Shown in **Figure 1(b)** in the upper row, a custom aluminum metal plate, with drawings detailed in the bottom row in **Suppl. Figure 1**, was fabricated in-house by the MPI SusMat workshop, which was placed into a TFS plastic loading bowl (part of the normal TFS life science workflow) referred to as “LN2 bowl.”

It has three pre-cut positions for fixing TEM grid boxes, two positions for loading a custom TEM holder^15^, top row in **Suppl. Figure 1**, as well as the golden base plate for loading the shuttle. The gold-plated metal plate has capability to host APT specimen holders (i.e. pucks), copper pre-tilt shuttles, etc. (Ferrovac AG, Zürich, Switzerland).

The custom copper shuttle has screw holes, including one where a SEM-clip is mounted to hold a clipped TEM grid in place parallel with the pre-tilt surface for EBSD. A second notch was cut into the pre-tilt face to fix a custom TEM holder in position, such that the holder with the grid could be positioned -20° off the electron beam optical axis, which is required to perform TKD, so that the electrons pass through the lamella and exit in the back to be collected by the EBSD detector. This geometry is different compared to conventional EBSD, where the sample surface to be analyzed is tilted to 70° with respect to the electron beam optical axis. The custom TEM grid holder is shown containing a clipped grid in **Figure 1(b)**.

After mounting the grids, the LN2 bowl was filled with LN2 and placed into the large antechamber or airlock on the nitrogen-filled glovebox (Sylatech GB-1200E, Sylatech GmbH, Walzbachtel, Germany), cycled three times quickly from 200 mBar to 750 mBar to avoid LN2 evaporation, and transferred into the glovebox as in the lower row of **Figure 1(b)**. Transfer under LN2 into an inert gas filled glovebox, as well as the use of appropriate cryogenically cooled UHV suitcase ^42,43,46^ avoids exposing samples to air between storage and instruments, as frost formation can destroy samples and sample cracking can occur as in **Suppl. Figure 2**.

The next steps of the workflow have been well documented for similar works ^45,47,48^. The gold-plated metal plate was then picked up from the LN2 bowl, as in the lower row of **Figure 1(b)**, and transferred to the LN2 bath, where it sits upon a platform on a horizontally sliding rod in a full LN2 bath. That metal plate remains there on its slider while the pre-cooled vacuum chamber and elevator (Ferrovac AG) was vented inside the glovebox. Then the elevator was lowered, the horizontal rod was used to slide the platform and metal plate onto the elevator, and the elevator was retracted. The chamber was then closed and pumped down to below 1 x10^-7^ mBarr. After that, the gate valve between the LN2 pre-cooled ultrahigh vacuum (UHV) cryo-transfer suitcase (UHVCTS)(Ferrovac AG), called “suitcase”, was opened, and its wobble stick extended to securely insert the pre-tilted shuttle into the suitcase. Finally, the gate valve was closed and the suitcase was removed and transferred to the cryo-FIB, as documented in ^42,49,50^.

#### SEM/FIB

The cryo-SEM/FIB used for this work is a TFS Helios 5 CX Ga^+^ (TFS, Waltham, MA, USA) equipped with a freely rotating Aquilos 2 like cryogenic stage and cryogenic EZ-Lift micromanipulator, was used to prepare specimens. Stage N_2_ flow was set to 190 mg/s with the stage heater temperature set to -190 °C for both stage and manipulator channels – the micromanipulator typically reaches -175 °C. The system was given thirty minutes to stabilize after cooling, following protocols in ^49^.

#### In situ diffraction by EBSD & TKD

For EBSD / TKD, the microscope has an EDAX Clarity Plus DED (EDAX/Gatan, Pleasanton CA, USA), which is a four chip TimePix-based detector, equipped also with a forward scatter detector (FSD). When the system was used in EBSD mode, the grid was placed under the SEM-clip and stage tilted 41° so that combined with the 29° pre-tilt holder, the sample was at 70° tilt, and the DED was extended into the chamber as shown in **Figure 1(c)**.

When TKD was performed, the custom TEM grid holder was loaded and the stage tilted to 23°, so that the grid was physically 20° negatively tilted (e.g. -20°) from the electron beam path / optical axis, and the DED was inserted so that a part of the scattered electrons (about those scattered between 30° to 90° from the primary electron beam) would reach the detector, as shown in **Figure 1(c)**. Schematically the difference between the EBSD and TKD procedures is shown in **Figure 1(c)**, e.g. the diffracted beam is sent to the detector from the surface versus transmitting through the sample.

Data was acquired using the EDAX APEX 3.0.6 software, at 10 kV and 1.4 nA unless otherwise specified. The working distance was 12 mm for optimal EBSD data, and 3.9 mm for TKD. The collected patterns were analyzed off-line in EDAX OIM Analysis™ 9.1 using the spherical indexing method with bandwidth 255 against a dynamical reference pattern for hexagonal ice (Ih) ^51–54^. To compensate for large variations in initial EBSD pattern intensity and contrast, additional image processing steps were applied. First, normally a mask (“Clarity”) was applied to eliminate boundaries between the four quadrants of the DED, typically with a width of 1 or 2 if required. Secondly, neighbor level parameter (NLPAR) pattern averaging was applied, using a search radius of 1.5 times the step size for nearest neighbor correlation.

An average background was reconstructed from the recorded patterns and subtracted from each pattern dynamically, followed by a quad equalization to correct for small variations between the individual sensors of the EBSD detector. If necessary, a static background was applied as well. Automatic brightness and contrast (ABC) were adjusted for each pattern image. Finally, the images were stretched to almost black and white with adaptive histogram equalization (AHE) to minimize projection gradients. With spherical indexing, the projection distortion as seen by the EBSD detector is removed and the patterns are fitted against a simulated spherical reference pattern that contains all possible diffraction bands. Spherical indexing is generally robust against both detector-and sample-induced distortions, such as pattern shadowing, detector intensity variations and tilt inaccuracies, projection center offsets, and specimen artifacts like holes or curtaining in FIB-produced lamellae. This robustness operates so long as the appropriate dynamic and static backgrounds corrections are applied, and masks are used to exclude artifacts when necessary ^52,55,56^.

The quality of indexing to assess the most likely orientation within those calculated by the software is expressed in a spherical Correlation Index (CI)^56^. Unless otherwise noted (e.g. for TKD), a spherical correlation index (CI) cutoff of 0.18 was used for the displayed or overlaid inverse pole figures (IPFs).

### Cryo-TEM lamella preparation and imaging

For this step in the workflow in **Figure 1(d)**, on-grid cryo-TEM lamellas were prepared following on-grid lamellas procedure commonly used in the life science community ^13,16,57,58^. Two parallel stress relief cuts are made, a few microns away from the targeted region-of-interest. These are significantly longer than the cut area. Top and bottom areas are milled away, and then progressively finer currents, ending in 1 pA and 30 kV, are used to polish the final lamella to a thickness of around 200 nm. A Cr lamella was pre-mounted on the micromanipulator at room temperature, and the liftout was then performed under cryogenic conditions, as described in Ref. ^59^.

For the last workflow step in **Figure 1(d)**, grids were taken for cryo-EM screening to verify lamella quality. These grids were taken to another facility in a LN2 transfer dewar and overview TEM images were collected on a Thermo Fisher Tecnai Arctica 200 kV cryo TEM with a Falcon 3 Camera.

### APT specimen preparation and analysis

Cryogenic lift-out (Cryo-LO) for APT was performed following the protocol described in Ref.^59^. A lamella of 99.9 % pure Cr (15×5×3 µm) was first lifted out at room temperature and attached to the micromanipulator. The temperature setpoints for sample stage and micromanipulator were then set to -190 °C; the stage cools to the setpoint, but the temperature of the micromanipulator to -175 °C. Lift-out was performed using a series of lines patterns on the interface between Cr and the region of interest to create an attachment via re-deposition^60^. A similar approach is used to attached the lifted-out bar to a sample support (36-post APT silicon microtip coupon, Cameca), and Cr-*in situ* sputtering used to strengthen the junction between the two^49,61^. With the stage tilted to 52° to face the ion beam, annular milling was used to shape the specimen with decreasing acceleration voltage and beam currents down to 5 kV, 15 pA, for the final steps. APT data was acquired on a Cameca LEAP 5000 XR atom probe (Cameca Instruments Inc, Madison, WI, USA). The base temperature was-233 °C (50 K), the target detection rate was varied from 0.5% to 0.7%. The data was acquired in laser-pulsing mode, with a pulse energy from 35 pJ and a repetition rate of 200 kHz. Data reconstruction was performed using Cameca’s Integrated Visualization and Analysis Software (IVAS) 3.8.20.

## Results

### EBSD Results

Figure 2(a) is a secondary electron (SE) micrograph of a cryo-TEM grid carrying sea urchin sperm on the pre-tilt FIB/SEM holder with a clip, with an additional stage tilt to be almost parallel to the ion beam, i.e. approximately only 8° off the optical axis of the FIB. EBSD was performed and Figure 2(b) is a so-called pattern region of interest analysis system (PRIAS)^62,63^ image, in which the contrast is close to that of backscattered electron micrographs. The orange dashed lines overlay are a guide to the eye to indicate where the Cu grid bars are.

**Figure 2:**
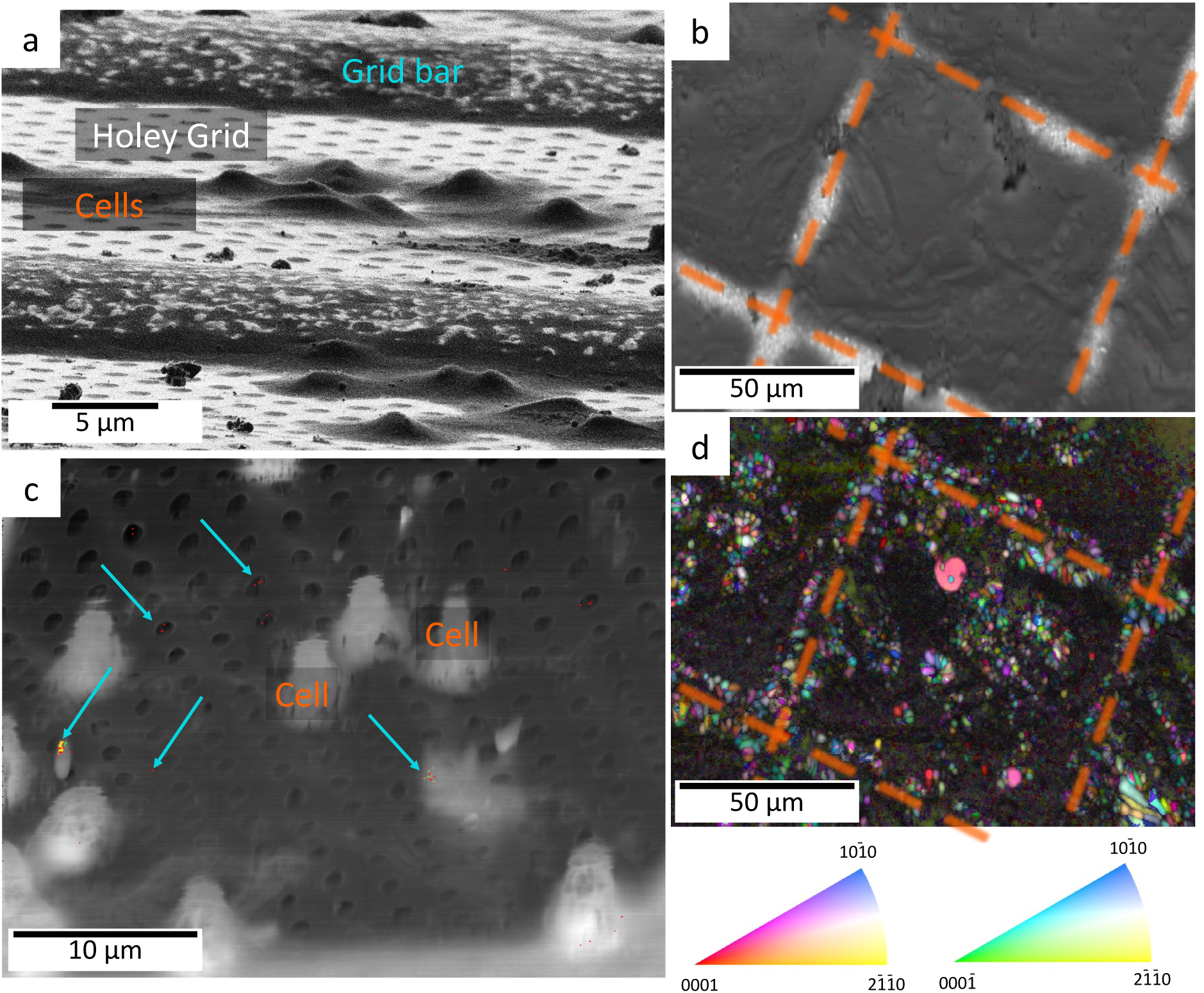
(a) Sea urchin sperm cells on holey Quantifoil grid. (b) PRIAS image, showing Cu grid bars with overlaid orange dashed lines as visual guides. (c) EBSD spherical correlation index mapping overlaid with grain orientation map, with small regions containing crystalline ice marked with blue arrows; note that the image was not tilt corrected, hence the visible distortions. (d) Crystal orientation map with respect to the scanning direction showing hexagonal ice, as well as partial ice on TEM Cu grid bars. IPFs color legends for Ih ice are shown.

In the CI map from the central part of a TEM grid square in Figure 2(c), regularly spaced holes from the carbon film on the grid are imaged in dark contrast, whereas cells appear in bright contrast, which suggests stronger scattering. Using spherically indexing and a CI threshold of 0.18, a grain orientation map was created and limited regions each containing only a few pixels and indicated by blue arrows, were readily identified as hexagonal ice (I_h_). Beyond I_h_ ice, the regions that do not generate signal and appearing dark are interpreted as vitreous. Another example is displayed in **Suppl. Figure 3**. On another grid, Figure 2(d) plots the grain orientation map overlaid onto the SEM-like DED image, evidencing significant crystalline ice on the Cu TEM grid bars, marked with orange dashed lines, where the cooling is locally not as fast and hence the risk of crystallization higher. The color code extends over two IPFs orientation triangles since hexagonal ice is not centrosymmetric, meaning that the atomic structure does not look identical when observed along or opposite to the c direction of the crystal ([0001] or [0001]) ^64^.

To further confirm that the indexing could distinguish the crystalline and amorphous regions of the sample, Figure 3 plots an EBSD pattern quality (PQ) map in gray scale obtained from a 70° tilted ice film. PQ Is equivalent to image quality (IQ) in the OIM software. Superimposed is a semi-transparent IPF. The underlying Cu grid is visible by a slightly higher PQ and marked by red dashed lines. Diffraction patterns obtained from four different regions, numbered 1 – 4 are also provided. (1) and (3) are from amorphous areas or areas with extremely fine-grained crystals. (2) is from a large hexagonal ice crystallite with good pattern quality. (4) is from a very small crystallite that can still be indexed by the spherical indexing process. Only hexagonal ice is found for the crystalline phase. Another similar example is shown in **Suppl. Figure 4**.

**Figure 3:**
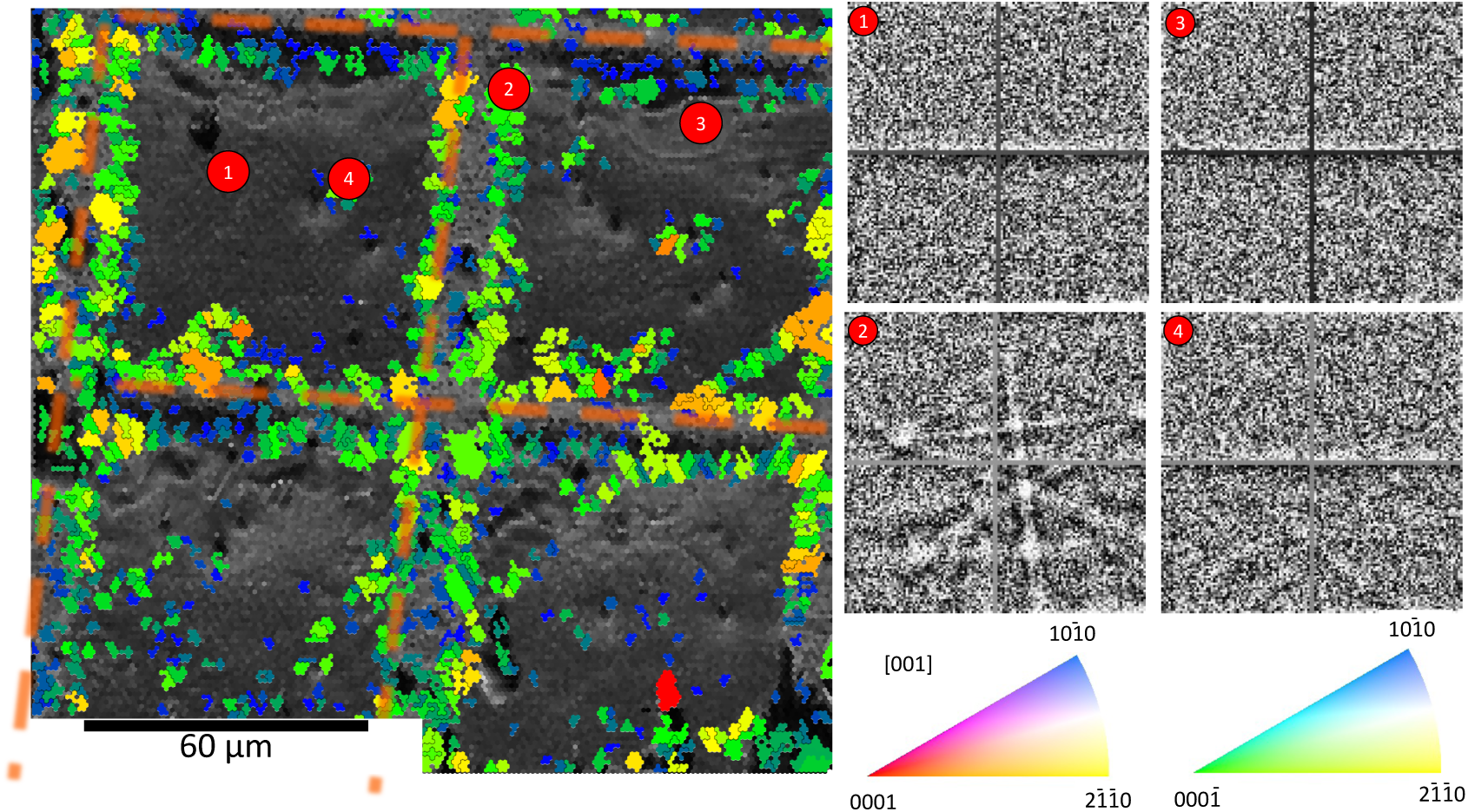
IPF superimposed to a grey-scale image quality map showing multiple grid squares analyzed by EBSD along with selected patterns from across the map as numbered 1 – 4. The overlaid orange dashed lines are visual guides indicating the location of the Cu grid bars.

### TKD

Previous work had suggested that TKD could be useful to validate lamella or grid crystallinity ^29^, where only one point of a defocused beam was used to acquire one Kikuchi pattern for a lamella. Here, after successful lamellae preparation, off-axis TKD was performed at an acceleration voltage of 10 kV, with beam currents from 700 pA. The polished lamella in Figure 4(a) showed no defects prior to TKD scans, whereas damage is readily visible in Figure 4(b), in the form of an induced gas bubble or similarly shaped artifact formed in the top portion and an array of holes at each scan point on the bottom side. Similar artifacts were imaged in other lamellas (Figure 4 **(c)** and Figure 4 **(d)**), indicating too high an electron dose. More severe damage were observed for higher acceleration voltages and currents. These lamellas were structurally damaged, rendering them unusable. Grid squares were also scanned, but regular partial shadowing by Cu grid bars resulted in limitations of the field-of-view, low quality of the indexing and only low-intensity Cu signals **Suppl. Figure 5 (a)**. No indexable patterns could generally be obtained from the thin ice films, but rather only from weak electron reflections from the Cu grid bars. Therefore, only patterns from the Cu grid bars were transmitted through the ice as in **Suppl. Figure 5 (b)**.

**Figure 4:**
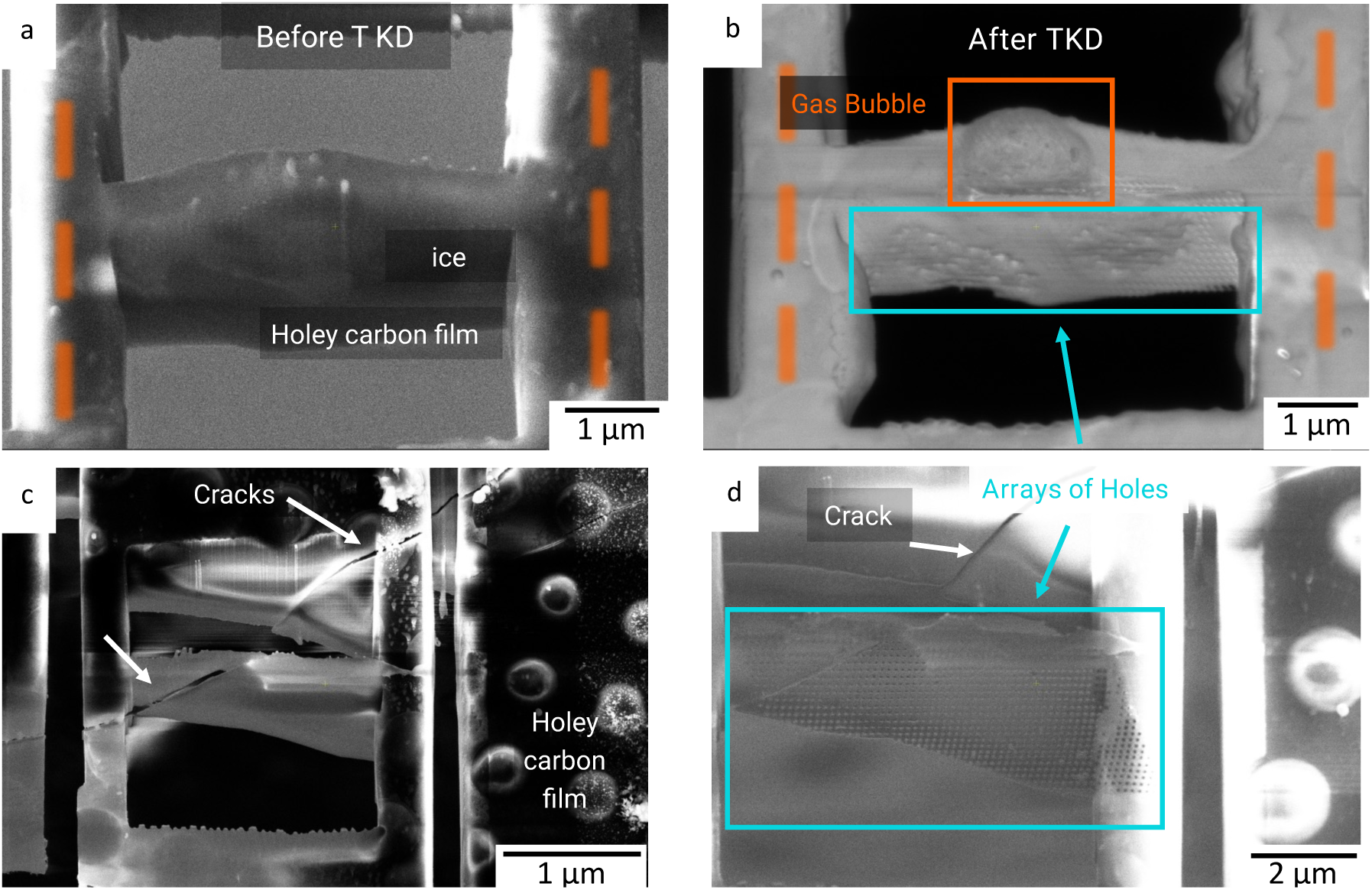
TKD results for lamellae (a) example ice lamella before TKD, containing the ice layer on top of the holey carbon film. The orange dashed lines are visual guides to indicate the location of the Cu grid bars; (b) same lamella after TKD, showing induced artefacts in the form of an apparent gas bubble along with an array of holes created by each scan data point; (c) ice lamella with a crack and (d)another lamella after TKD scan, showing array of holes at the TKD sample intervals.

For light, weakly scattering materials like water ice, particularly for a 100 – 200 nm thick layer on a plunge-frozen cryo-TEM grid, the relative mean free electron path and the interaction cross-section are expected to be low. A Monte-Carlo ^65^ electron trajectory simulation was performed and is mapped in **Suppl. Figure 6**. This shows that most of the backscatter electron signal for EBSD, i.e. those with the lower energy loss, is near 200 nm in diameter right in the subsurface area. Although these simulations do not consider “thermal diffuse scattering” that is responsible for Kikuchi diffraction, they provide an approximate idea of the interaction volume. In normal incidence, this volume would be reduced down to roughly the thickness of the lamellae imaged in Figure 4, which suggests that sufficient scattering to generate detectable TKD signals does not occur. Similar problems have been found for pure Li for instance, where a thickness of more than 1 µm was found necessary^66^.

### Cryo-TEM lamella preparation and imaging

An example of a cryo-TEM lamella preparation process is shown in Figure 5(a), with the stage tilted as explained in the Methods section, which explains that only a part of the image is in focus. The stress relief cuts help avoid cracking of the ice once the main cuts are made on either side of the sample of interest. Since the cells were at least a micron tall, the nearly lateral cuts at 7-8° off the FIB axis yielded lamellas several microns long with removal of perhaps a few hundred nanometers of material per side. This lamella, imaged in the SEM at 1 kV, 21 pA, Figure 5(b), contains visible cross-sections of cells. Note that the micrograph was obtained with a stage tilt, which explains why it is focused only on the cellular components within the lamella. Lamellae were unloaded from the SEM/FIB into the glovebox through the pre-cooled UHV suitcase, and transferred back into cryo-TEM grid boxes. Figure 5(c) is a top-view cryo-TEM image that demonstrates that the lamellas were intact and free of frost despite moving the grids across multiple facilities (> 50 km apart). If a cryo-TEM lamella is not properly handled and warms up even midly, the ice can sublimate and the resultant grid would have no remaining vitreous ice and a small amount of frost with a “freeze-dried” appearance as in **Suppl. Figure 2 (b)**. This demonstrates that the proposed workflow allows for handling biological samples, from end to end, under conditions that avoid ice crystalization that would result in damage to the sample of interest.

**Figure 5:**
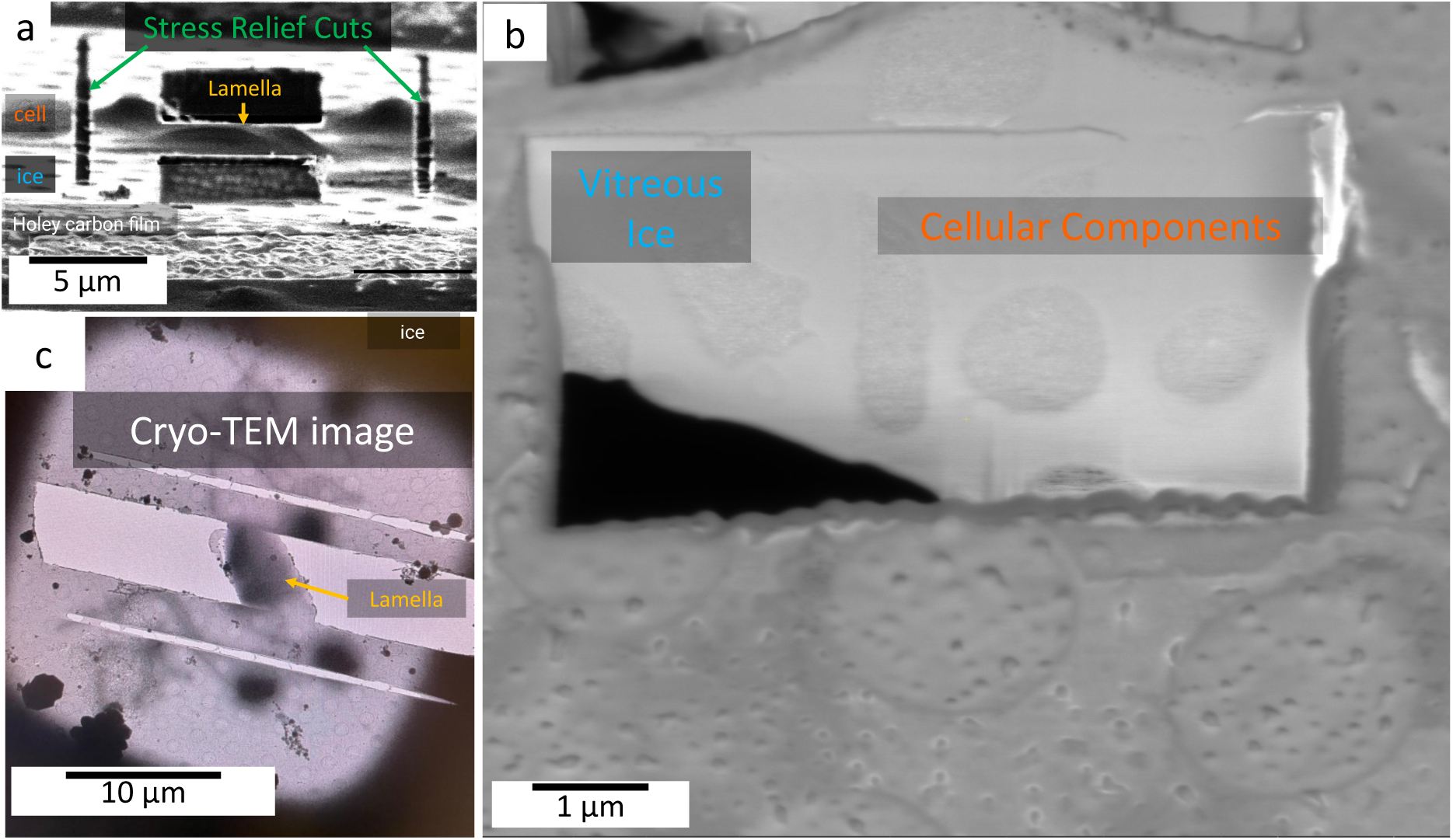
(a) partially thinned lamella from one cell (darker contrast), along with two stress relief cuts along the sides to prevent bending from thin film stress in the carbon membrane with thin ice. (b) polished final lamella, with an approximate thickness of 200 nm, showing various sea urchin sperm cross sections in secondary electrons inside the SEM/FIB. (c) cryo-TEM image of a similar but not identical lamella showing intact lamella with no visible surface ice contamination, amenable for imaging and tomographic analysis.

## Discussion

The life science community has well-developed workflows for on-grid lamella preparation including extensive vendor-specific automation, as well as open source workflows like SerialFIB ^67^, as reviewed in ^68^. Likewise, life-science vendor-specific solutions (e.g. TFS Aquilos 2, etc.) exist and are used in higher-end facilities for automated specimen preparation with transfer into automated cryo-TEMs. However, the vitrification yield varies across users, as well as specimen and preparation method. That is problematic because ice crystals formation destroy native biological structures due to the volume expansion ^26^, and these crystals are typically only detected on the lamellas in the TEM, which is the second last step in the workflow. This limits the throughput and wastes microscope resources, time and budget. Therefore, it is highly desirable to assess the vitrification of samples at earlier steps of the workflow to decide if a given lamella is of sufficient quality for cryo-TEM imaging or analysis. Previous attempts to validate sample integrity in the FIB/SEM using STEM detectors^29^.

Here, we introduced the design of custom holders which allow reliable transfer of clipped cryo-TEM grids into a materials science FIB/SEM instrument, using a sensitive DED detector in a conventional EBSD configuration to assess the presence of crystalline ice within plunge frozen cryo-TEM grids carrying cells. We demonstrate proof of principle that EBSD using a DED can be used to show the presence of hexagonal water ice I_h_ on the surface of a grid, including on the biological sample of interest. In contrast, areas that produce no diffraction signal are assumed to be composed of vitreous ice. This information is very valuable to the operator since specimen preparation will only be performed on the vitreous areas of the grid. That means that the time and resource consuming sample preparation can be avoided on crystalline areas of the sample which makes the overall workflow a lot more efficient.

This demonstration opens the door to using detectors already present on numerous material science cryo-FIBs, in facilities equipped with correlative microscopy toolsets and transfer equipment, to assess the if samples are vitreous or not before preparing lamellae for cryo-TEM or specimens for cryo-APT. We introduced here additional, simple custom fixturing which allow reliable transfer of cryo-TEM clipped grids made either on-site or in another facility into materials science laboratory workflows. This loading procedure uses standard equipment in existing specialist labs for cryogenic vacuum transfer for correlative microscopy and is compatible with the dovetail latching mechanism used in typical shuttle systems as seen in Figure 1(b). Clipped cryo-TEM grids were reliably moved into the glovebox and back out, without devitrification or additional surface ice contamination, providing useful lamellae.

### EBSD-induced crystallization and damage

Proving that a material is crystalline based on EBSD alone can be done when diffraction patterns are recorded, enabling their indexing. Here, the regions that do not yield indexable diffraction patterns are assumed to be amorphous. Multiple reasons exist that could prevent diffraction patterns to form, including for instance that the ice is made to grains too small for the large volume probed in this very light material, i.e. in which scattering is limited, and patterns are too weak to be indexed. As de Winter et al. reported that electron-imaging can induce crystallization^29^, we performed comparative imaging of the same region before and after EBSD scans across a range of beam acceleration voltage and current.

Maintaining the acceleration at 10 kV, we have systematically varied the beam current in the range 0.7 – 2.8 nA, and the raster step size from 0.3 – 1 µm. Depending on the imaging conditions, morphological changes of the sample could be readily visible. Figure 6(a) is a secondary electron micrograph of a cryo-TEM grid carrying sea urchin sperm cells. Some cells are circled in green as a guide to the eye. Following an EBSD scan at 10kV, 0.69nA, and a 1.0 µm raster step size, the secondary electron micrograph of the same region is displayed in Figure 6(b) showing no visible changes. Figure 6(c) is the PRIAS image providing a view of the EBSD detector, and the corresponding EBSD crystal orientation map is plotted in Figure 6(d) that confirms the presence of ice crystals in some regions of the sample, probably related to frosting and contamination during the transfer into the glovebox, but no diffraction patterns could be indexed across most of the sample area.

**Figure 6:**
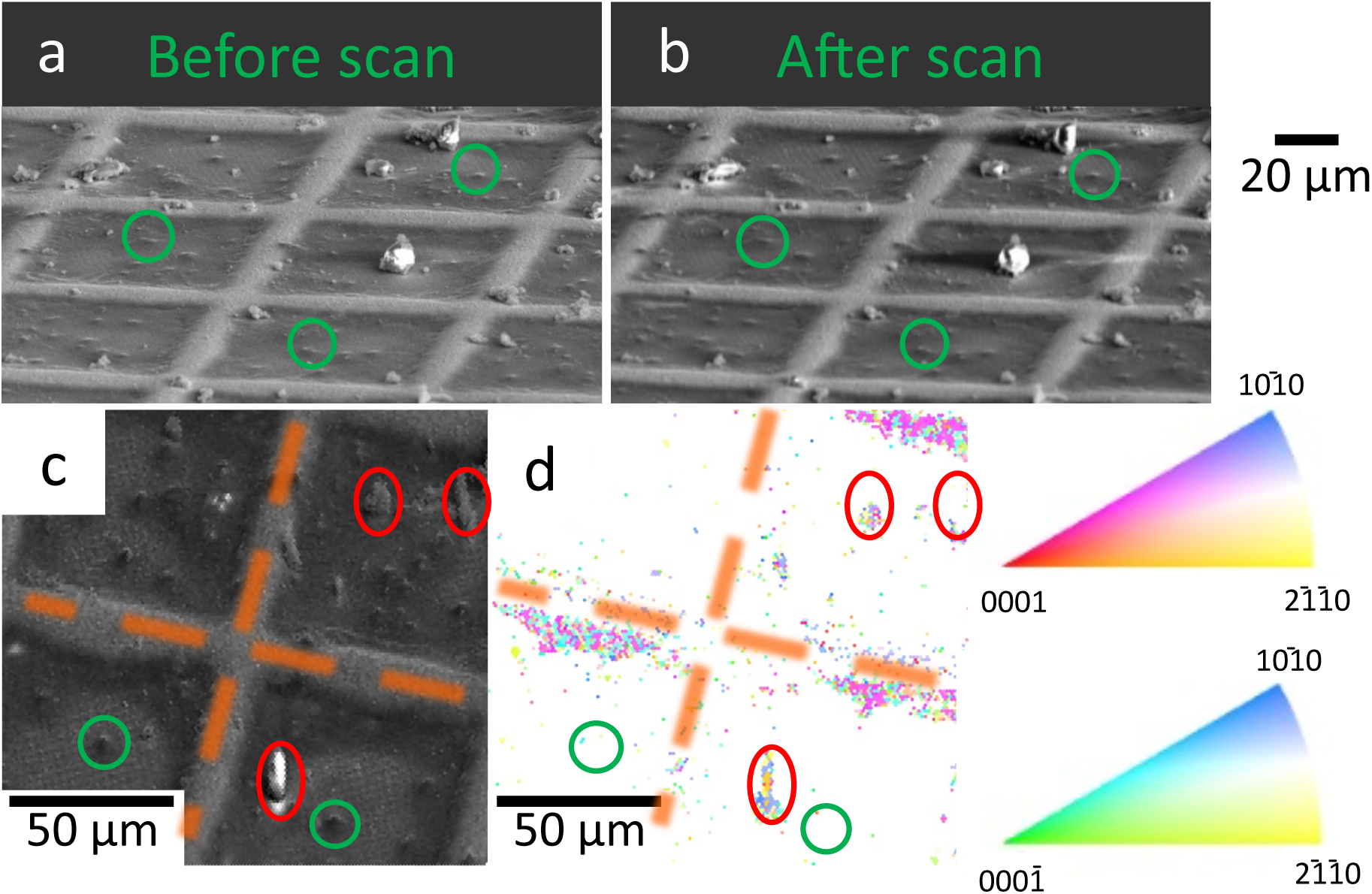
secondary electron micrograph of a cryo-TEM-grid carrying sea urchin sperm cells (a) before and (b) after an EBSD scan at 10kV, 0.69nA, and a 1.0 µm raster step size; corresponding (c) PRIAS image and (d) IPF from the EBSD scan indicating the presence of some ice crystals from frost contamination indicated by red ellipses, whereas cells delineated by green circles do not appear.

Figure 7 mirrors Figure 6 for a higher beam current of 2.8 nA and a shorter raster step size of 0.3 µm. The comparison of the electron micrographs before and after scan in Figure 7(a) and Figure 7(b) evidences a change in morphology, with the apparent growth of ice crystals on the cells that are within the scanned area. This is further supported in the PRIAS image in Figure 7(c) and the corresponding crystal orientation map in Figure 7 **(d)** in which crystals are spatially correlated with the location of the cells and grid bars.

**Figure 7:**
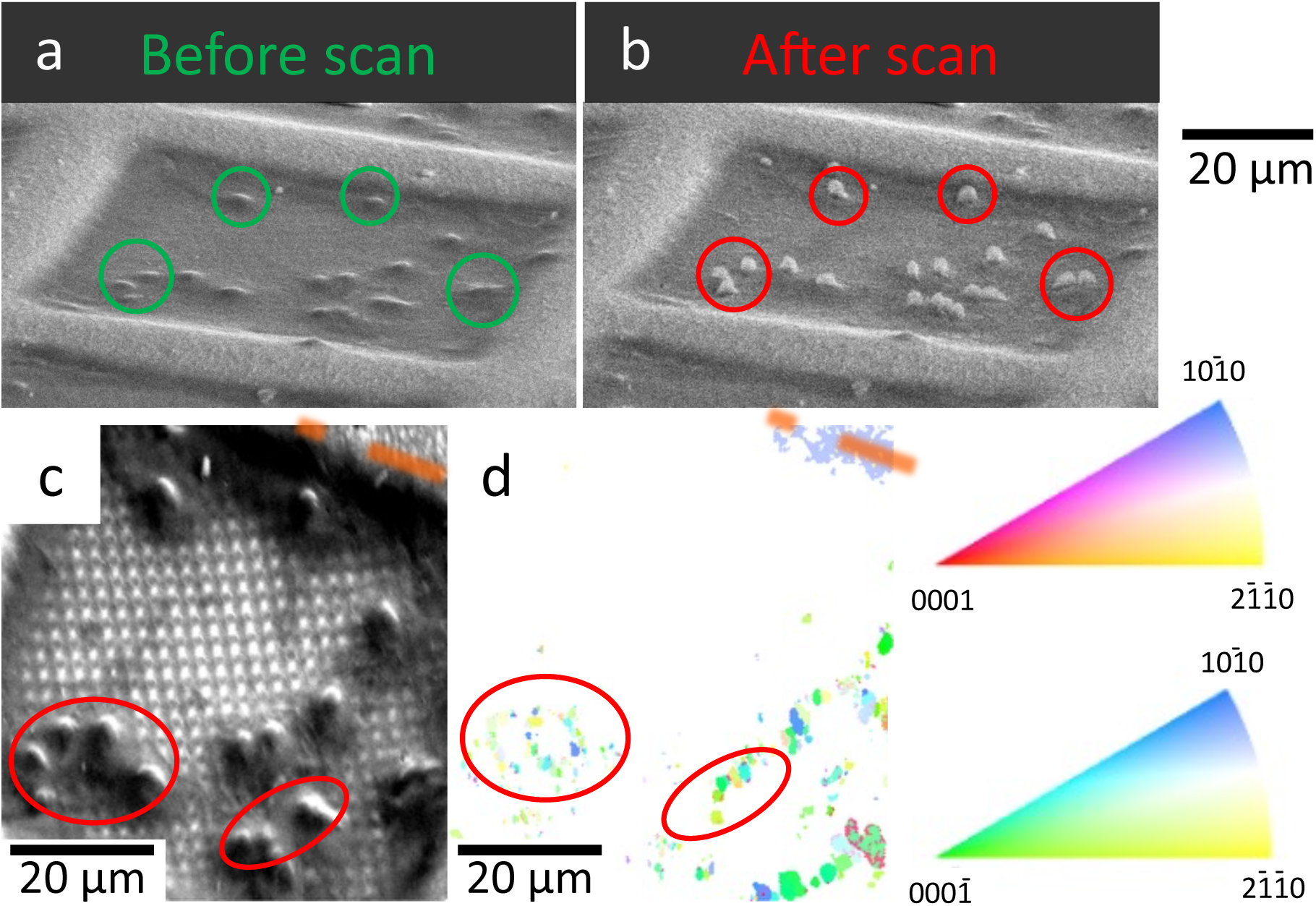
secondary electron micrograph of a cryo-TEM-grid carrying sea urchin sperm cells (a) before and (b) after an EBSD scan at 10kV, 2.8nA, and a 0.3 µm raster step size; corresponding (c) PRIAS image and (d) IPF from a second EBSD scan indicating the presence of some ice crystals, including directly on the cells of interest.

This exercise allowed us to first support our hypothesis that the areas of the sample that are not providing indexable patterns are amorphous; second, we could map satisfactory imaging conditions leading to sufficient signal and limited damage. The results of our systematic investigation of damage is summarized in Figure 8, in which the red points are combinations of current and step size with visible damage to the sample, primarily in the form of the crystallization of the ice as in Figure 7, and in green are combinations leading to no visible damage as in Figure 6. In between, hints of limited changes to the sample morphology could be seen, yet the corresponding EBSD pattern did not reveal crystalline ice, which could simply be due to insufficient signal. The shaded regions should be seen as indicative.

**Figure 8:**
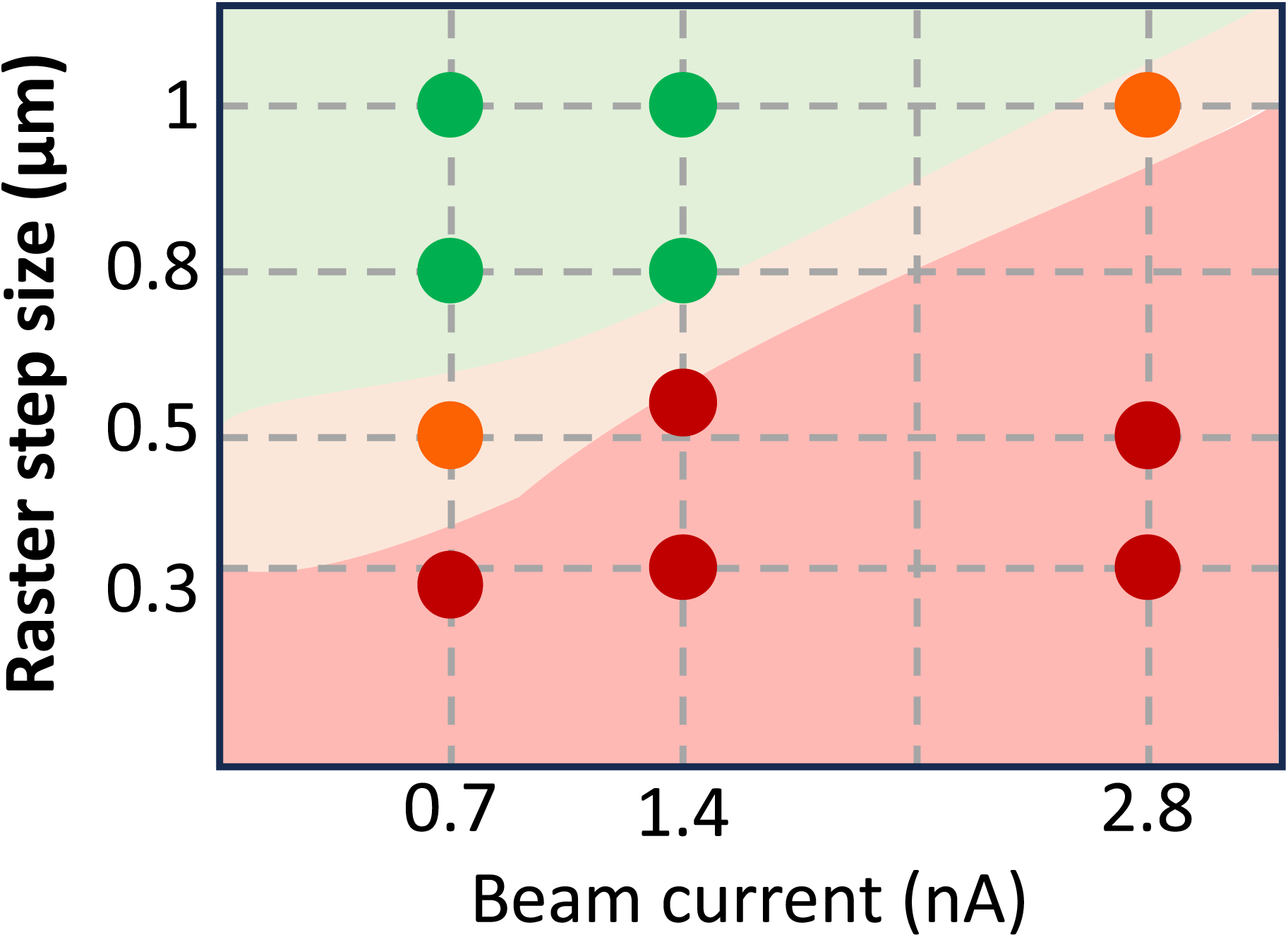
visible damage from the EBSD scan for a range of beam currents and raster step size, with red corresponding to measurable crystallization by EBSD, orange is damage only visible from SEM and no crystal formation in EBSD, and green reflects combinations leading to no visible crystallization or morphological changes of the sample by either of the two techniques. The shaded regions are a guide to the eye.

As first established in the cryo-TEM field, the electron beam damages ice samples, even at cryogenic temperatures ^34,69,70^. Cryogenic temperatures slow such damage, but do not stop it, vis several mechanisms ^69,71,72^. Vitreous or amorphous ice actually comes in different densities, where the electron beam can induce shifts between them^73^, as well as devitrify them if heated ^74^. In the lower energy regime of the SEM, the electron beam can induce sputtering, sublimation, and ice loss, as well as gas bubble formation ^75,76,77,78^. Therefore, dose-dependent damage takes place and must be carefully considered ^79^.

### Perspective on SEM-based ED for cryo-TEM screening

SEM-based ED can only evidence crystalline area where there is sufficient signal present to assess crystallinity. The implementation of ED that were used herein cannot generate Thon rings, like a standard TEM diffractogram, to differentiate amorphous from crystalline areas of a sample. Although it is extremely difficult to prove a negative, in that amorphous ice will not diffract and hence not generate signal in principle, realistically, the assumptions regarding crystallinity if insufficient signal or signal quality at a given pixel or feature appear reasonable. As we have demonstrated, particularly for off-axis TKD, there are practical considerations that can preclude the use of SEM-based ED for assessing whole-grid ice quality and crystallinity, including ice thickness due to poor scattering (**Suppl. Figure 6**), scattering from the metals from the grid and the clipped ring that can obscure the low-intensity signal from the ice (**Suppl. Figure 5**).

We have demonstrated that EBSD can be used to assess the presence of hexagonal ice which is expected for surface contamination and improper vitrification of plunge frozen biological samples. We optimized our data acquisition and indexing parameters and could detect meaningful diffraction patterns on specific locations of the grid. The presence of crystalline ice is expected from biological specimen preparation using common vitrification procedures producing cryo-TEM grids that suffer from local ice contamination and failed vitrification. Another critical aspect for implementation withing cryo-TEM workflows is the field of view possible within a reasonable time. Examples of EBSD map on multiple 2×2 grid squares are presented in Suppl. Figure 9. These three fast scans were performed within less than 20 min, and the spherical reindexing took ca. 3 min on a workstation. Integrating in-situ ED in the SEM as part of cryo-TEM specimen preparation workflows hence appears viable.

Multiple microscope and detector configurations are possible leading to differences in the efficiency of diffraction, i.e. the conversion between the dose of incident electrons onto the sample and diffracted electrons reaching the detector. Herein we could show that there is a higher chance of thermal diffuse scattering in the volume for a certain total number of incident electrons in an EBSD configuration, compared to TKD, and even if the electron’s energy is spread, there is sufficient signal to be detected by the highly sensitive DED and indexed. The configuration of annular DEDs located around the pole piece of the SEM allows also for reflection Kikuchi diffraction (RKD) ^80^, which was not tried here but could be performed with the grid lying flat on the stage, potentially facilitating further specimen preparation steps. On-axis TKD could also be revisited^29^ and optimized, in the hope that this could help further reduce the damage, particularly through the use of low-dose with a defocused beam, low voltage, and on-axis transmission scattering. The interaction volume for such a configuration is evidenced in the Monte-Carlo ^65^ electron trajectory simulation **Suppl. Figure 10**, highlighting the differences with the grazing incidence in **Suppl. Figure 6**, and indicating that each combination of microscope and detector would need optimization for accelerating the collection of high-quality patterns, while lowering the electron dose and acceleration voltage to minimize the risk of damage to the sample ^80^.

### Potential for correlative cryo-TEM-APT

Cryo-TEM yields mainly structural information for molecules and not the chemical identity of each individual atom. Since biomolecules are very sensitive to radiation damage, typically only very low electron doses are used to image them^36^, and even cryogenically cooled, vitrified specimens are normally not analyzed by using energy-dispersive X-ray spectroscopy (EDX) or electron-energy-loss spectroscopy (EELS), although recently proof-of-concept EDX in cryo-TEM has been used to identify light ions ^81^. In the materials sciences, atom probe tomography (APT)^82^, a form of mass spectrometry (MS), is often used to complement SEM and TEM to provide nanoscale compositional information in 3D ^46,83^. Identical SEM/FIB workflows are used for APT site-specific specimen preparation^84^ facilitate direct correlative imaging of the same microstructural feature by the different techniques ^85,86^.

These specimen preparation workflows were recently expanded to cryogenic temperatures ^59,60^ along with by UVH transfer protocols^87^. These have enabled analysis of frozen liquids^88,89^ including aqueous solutions ^50,90–93^ as well as other . Despite long discussions of APT’s potential for biology^94,95^, limited data has been reported for biomolecules in their native environment^90^, and processing of the data remains challenging^96^. Assessment of specimen crystallinity before site-specific preparation is likewise crucial to avoid damaged regions^97^. The geometry of a clipped TEM grid along with the tilt of the stage make the lift-out process particularly challenging. As demonstrated in **Suppl. Figure 11**, a lamella was successfully lifted out via redeposition welding^49^ from a grid carrying the sea urchin sperm cells, and sliced into 2–3 µm chunks. Each slice is then sharpened into a needle-shaped specimens, as in Figure 9(a), and the whole support is finally transferred from the FIB into the atom probe through the cryo-UHV suitcase. In this proof-of-principle experiment, two cryo-UHV suitcases were used, one for the APT coupon and one for the cryo-TEM grid. In the future, new cryo-stage top-plate designs or sample holders that can hold multiple shuttles may solve this problem and allow for both the sample support and the TEM grid holder to be present in the cryo-FIB chamber simultaneously.

**Figure 9:**
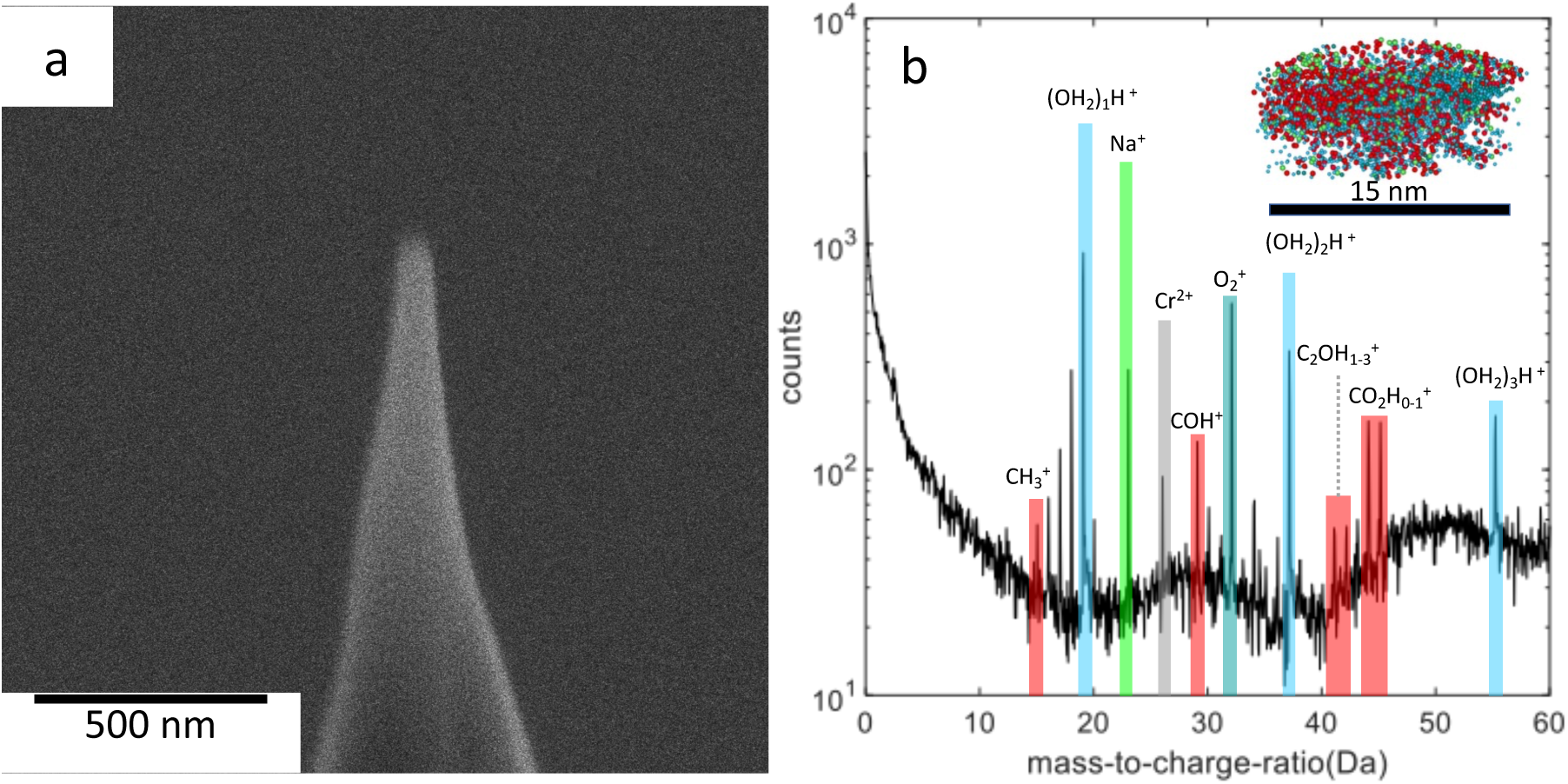
(a) sharpened APT specimen following annular milling; (b) resulting mass spectrum from the limited preliminary data.

Only limited data could be acquired here before sample fracture. Figure 9(b) plots a mass spectrum from one of the short acquired data sets that contains the protonated water-based molecular ions, (H_2_O)_n_H^+^, that are typically detected in the analysis of water ^93,98,99^. Additionally, peaks are detected that can be associated to Na^+^ ions from the saline aqueous solution ^100^, and a range of C-and O-containing fragments found in the analysis of organic matter by APT^96,97,101^, all of which can be attributed to the buffer solution. The Cr-ions are related to the redeposition welding process. This preliminary work opens the path for performing APT on plunge frozen samples on cryo-TEM grids and enable correlative workflows.

## Conclusions

We have introduced the design of new holders and jigs to facilitate transferring clipped cryo-TEM grids used in life science workflows into a cryogenic workflow initially designed for materials science. We have demonstrated EBSD with a DED to assess the crystallinity of ice on grids directly inside the FIB/SEM before further specimen preparation is undertaken. TKD in off-axis geometry proved unsuitable. We fabricated on-grid lamellae and transferred them back out through UHV suitcases and glovebox to image in a cryo-TEM without surface ice contamination. Despite additional steps for sample introduction and handling via the glovebox and UHV cryo-suitcases, the proposed workflow did not lead to additional contamination or structural changes to the sample. Our work paves the way for assessment of sample crystallinity readily in the SEM/FIB, skipping the conventional screening step in the cryo-TEM, along with correlative studies with e.g. APT. Future developments will facilitate automation of these workflows, validating realistic speeds for complete grids, as well as automating stage movement and establishing minimum speeds to assess crystallinity and screen the suitability of samples for high-resolution cryo-imaging and analysis.

## Acknowledgements and Funding

E.V.W., C.W. and B.G. thank the Deutsch Forschungsgemeinschaft (DFG) for their generous support through BG’s 2020 Leibniz Prize. T.M.S. gratefully acknowledges the financial support of the Walter Benjamin Program of the German Research Foundation (DFG) (Project No. 551061178). The authors thank Rainier Lück, Ralf Selbach, and the MPI SusMat machine shop for their help in improving TEM grid holder design and fabrication. The authors acknowledge the assistance of Katja Angenendt and Christian Bross in optimizing EBSD acquisition. The authors acknowledge Monika Gunkel and the StruBITEM facility at the University of Cologne for their assistance with cryo-TEM grid preparation.

## Data Availability

Data from this work is available from the corresponding authors upon reasonable request.

---

**Suppl. Figure 1:**
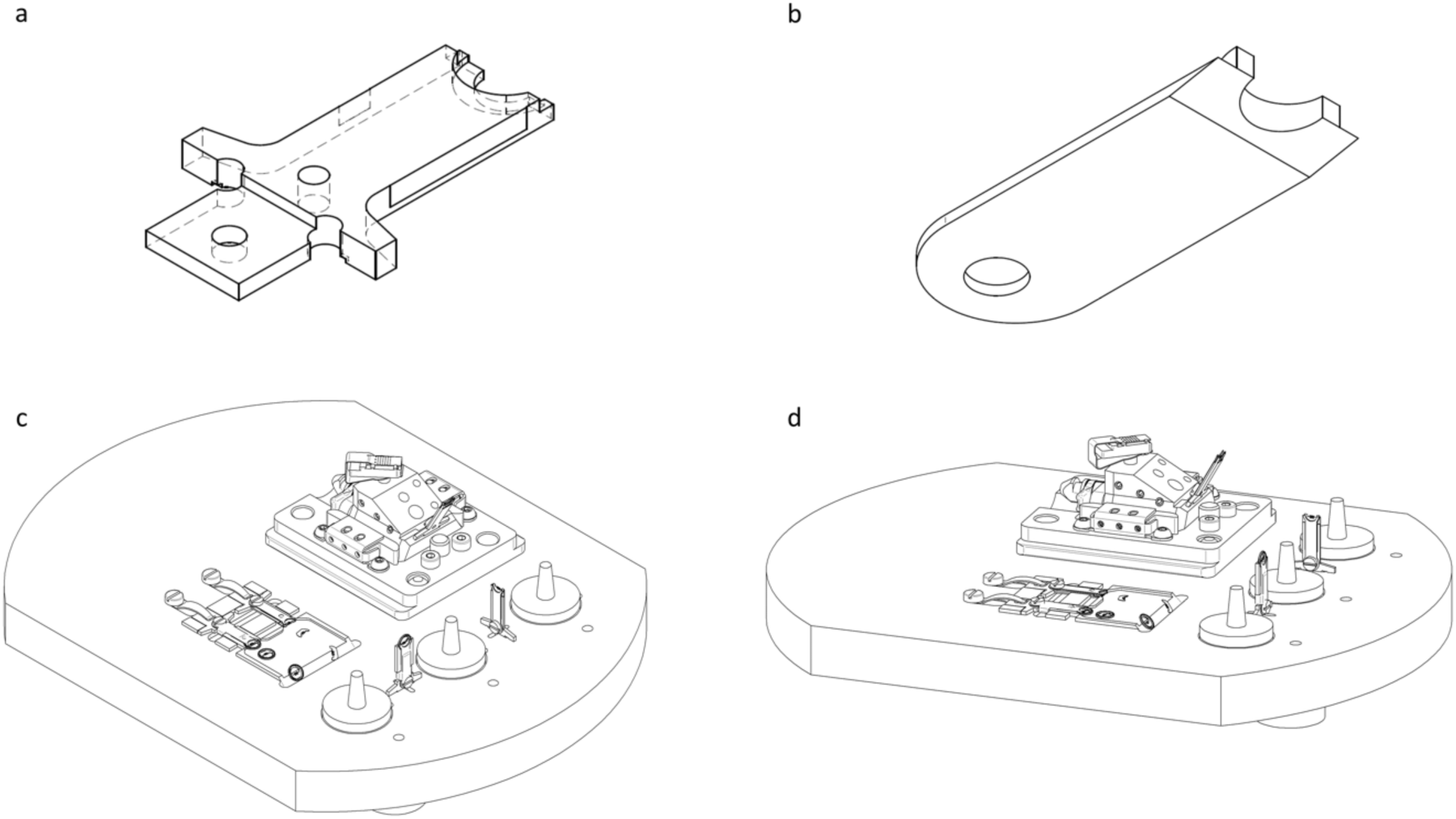
(a) bottom part of custom cryo-TEM clipped grid holder (b) top part (c) schematic view of metal loading plate, inclined side view (d) side view with different angle to better illustrate height differences.

**Suppl. Figure 2:**
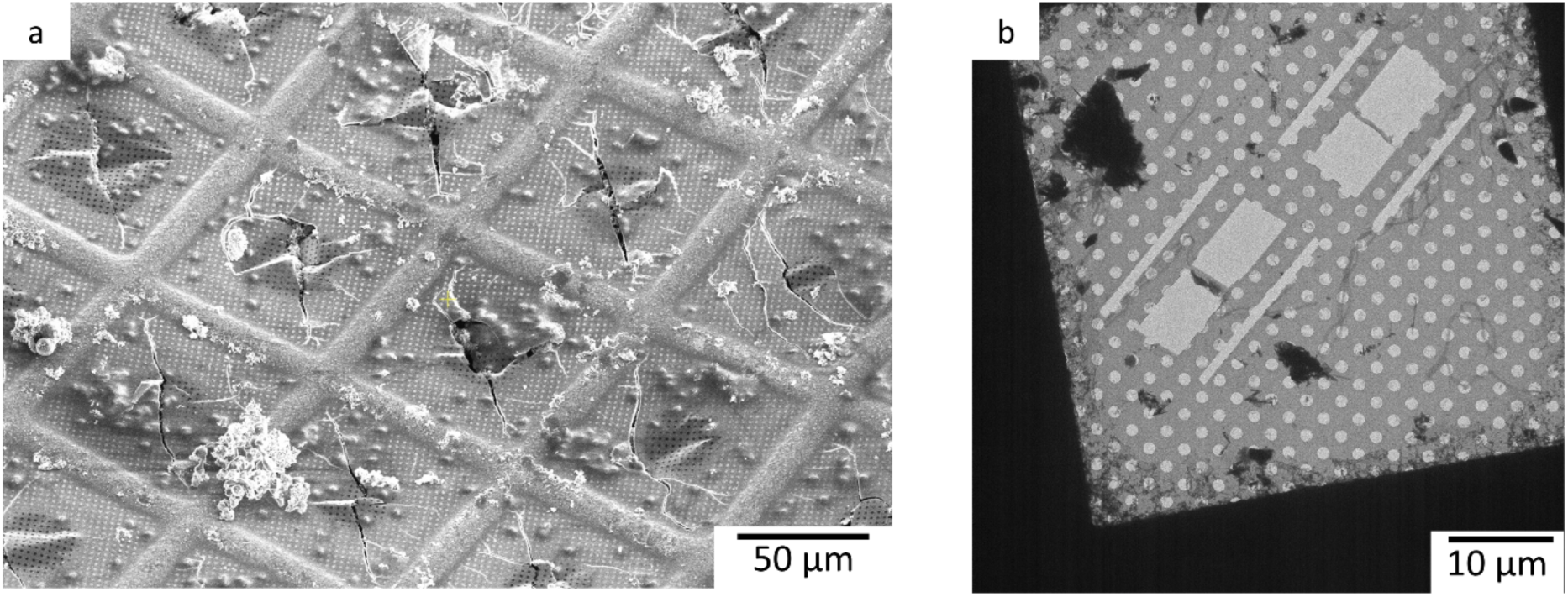
Example failure modes for cryo-TEM grids (a) surface ice (frost) contamination and ice cracking (b) sublimated (e.g. freeze-dried) grids with no remaining vitreous ice and a small amount of frost

**Suppl. Figure 3:**
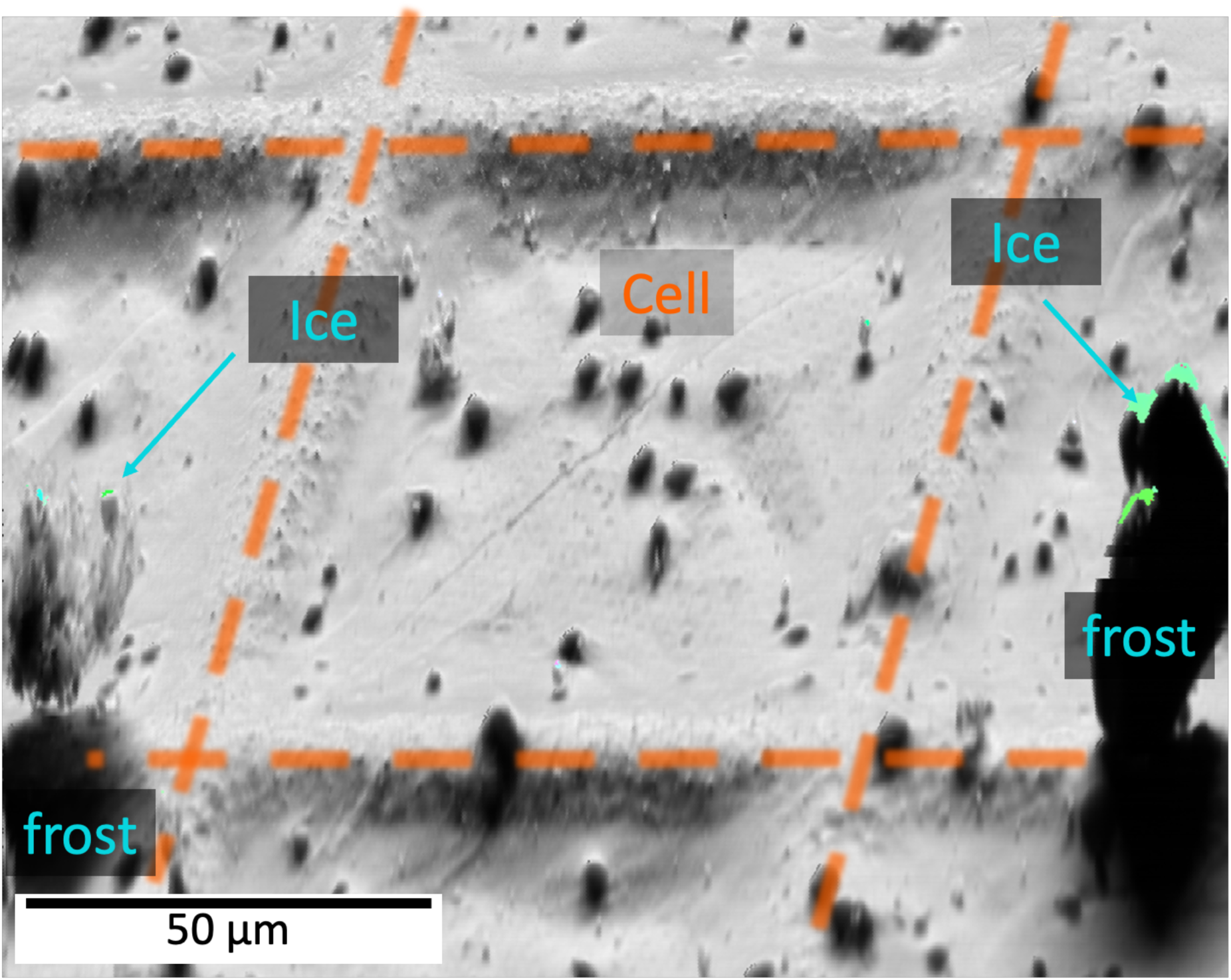
a DED image with a higher intensity of backscattered electrons. Crystalline regions are indicated by blue arrows. The surface ice crystals are up to tens of micrometers in size and block the diffracted electron beam, except at their edges. Cells form the topography in the middle of the main TEM grid square. Orange lines are overlaid as a visual guide to indicate the TEM grid bars. Note that the image was not tilt corrected, hence the visible distortions.

**Suppl. Figure 4:**
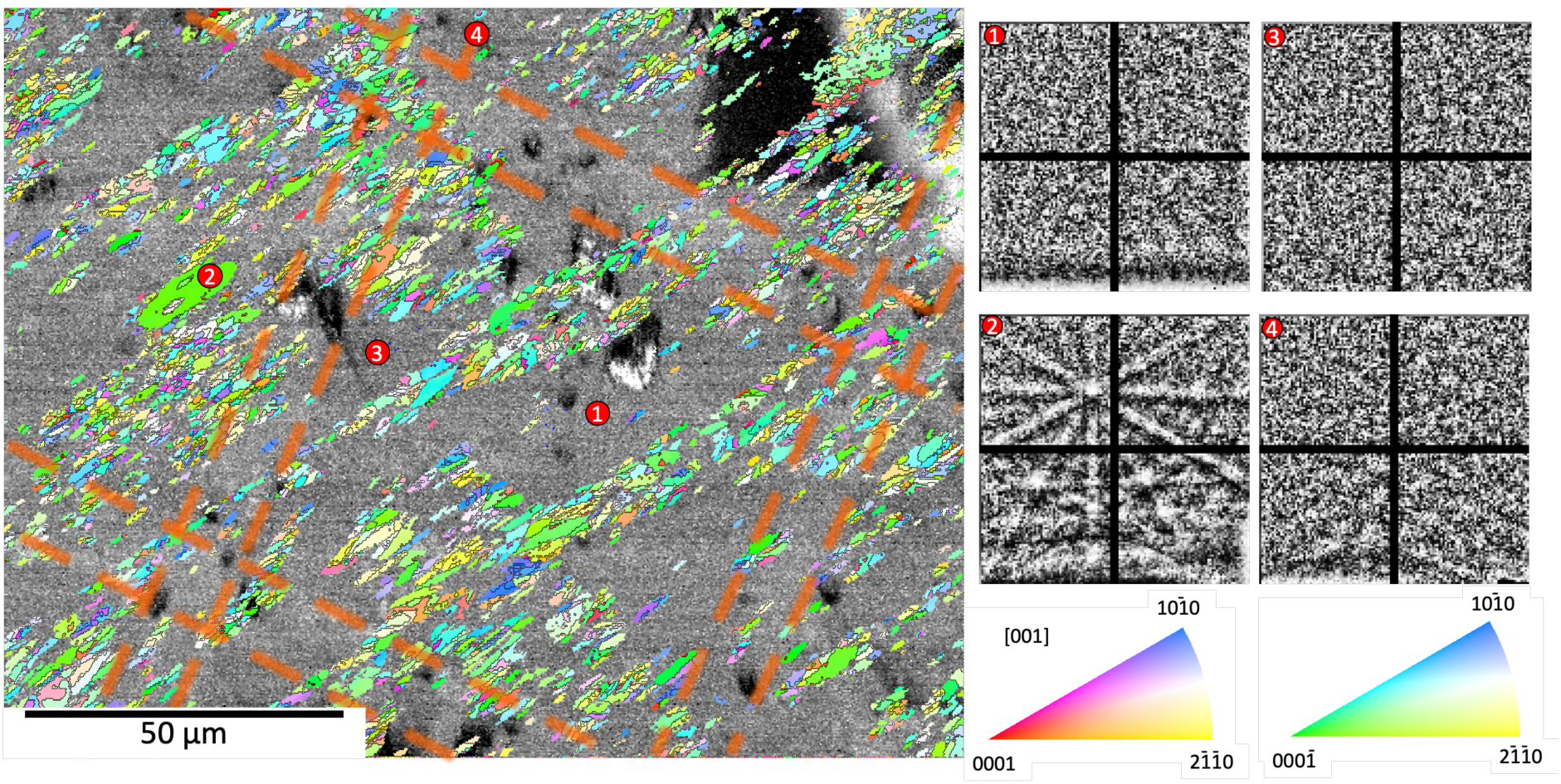
IPF superimposed to a grey-scale image quality map showing multiple grid squares analyzed by EBSD along with selected patterns from across the map as numbered 1 – 4. The overlaid orange dashed lines are visual guides indicating the location of the Cu grid bars

**Suppl. Figure 5:**
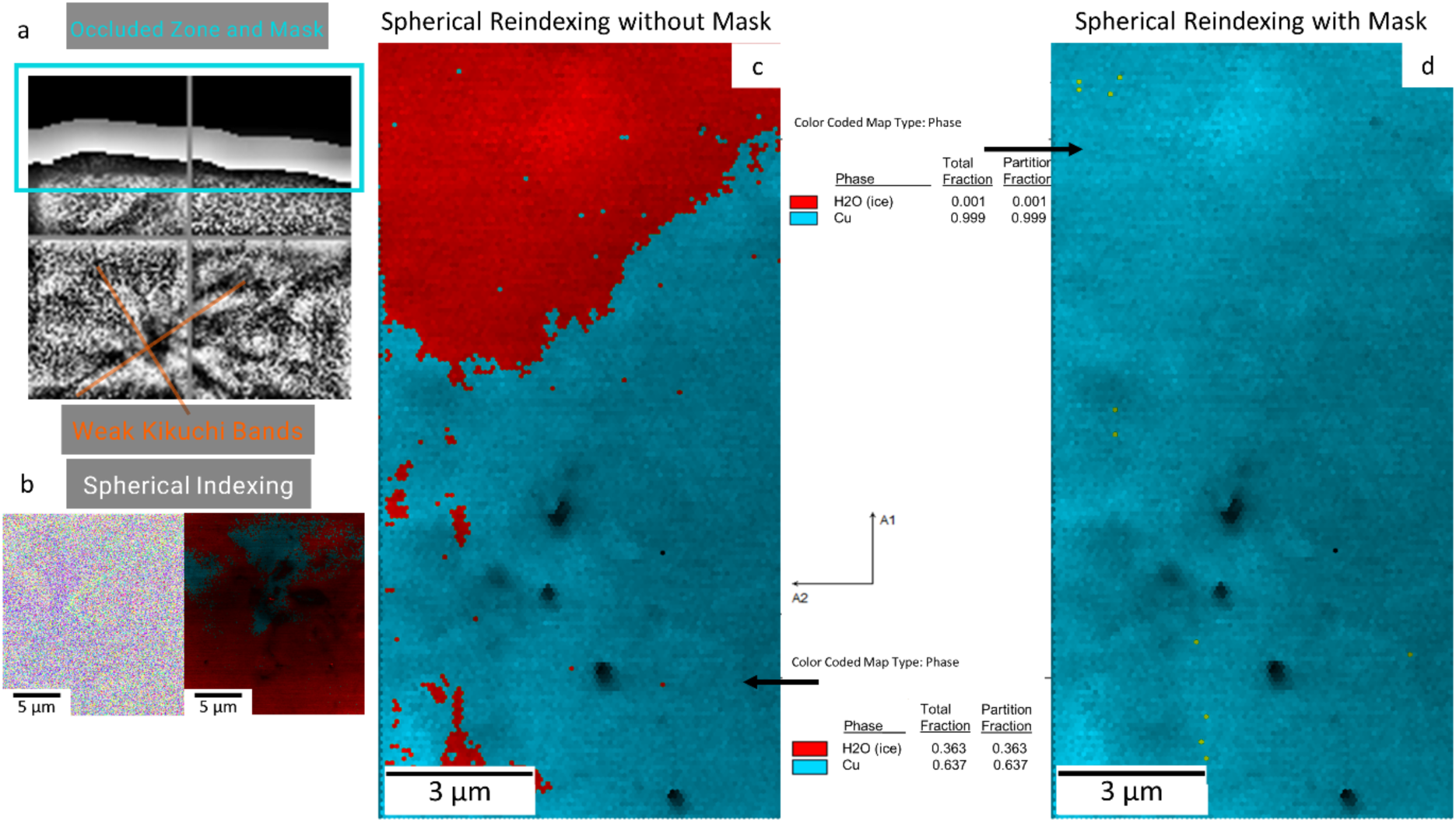
Grid TKD scan results (a) example TKD patterns taken by DED. Anything in the captured two-dimensional image of the pattern that breaks band symmetry through occlusion leads to indexing failure, such as the top area in (a). (b) initial results and spherical indexing refinement; occlusion limits the accuracy of the indexing, as shown in the left side, with low CI for the indexed area and results resembling random static. (c) phase identification using spherical indexing; the occluded area caused the pattern identification to have difficulty correctly phase matching. (d) phase identification using spherical indexing with mask; the term “mask” here means that a specific shape was applied to exclude the occluded area of the captured 2D image, as in the blue area marked in (a). When that area was excluded, more correct indexing result was obtained, but only signal from the Cu grid bars could be retrieved.

**Suppl. Figure 6:**
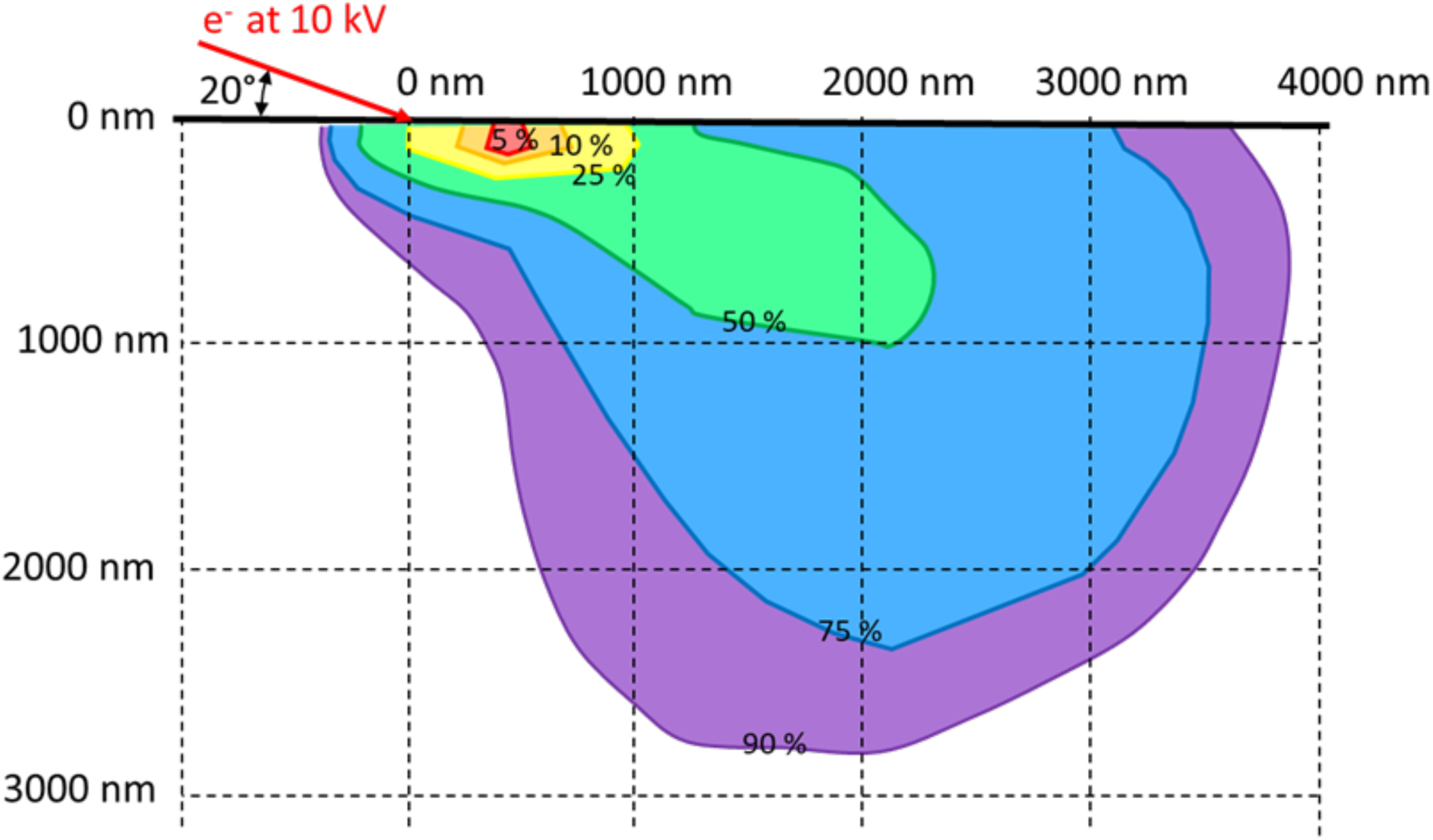
Casino Monte-Carlo trajectory simulation showing the total interaction volume for different energies with respect to the incident electron energy. The simulation is for 70° tilt (20° grazing incidence) at 10 kV acceleration voltage. The red region corresponds to the volume where electrons have almost no energy loss, i.e. from where originate most backscattered electrons. The area where the most significant amount of energy (blue) is introduced into the material is near 3 µm in width but confined in depth to near 2 µm.

**Suppl. Figure 7:**
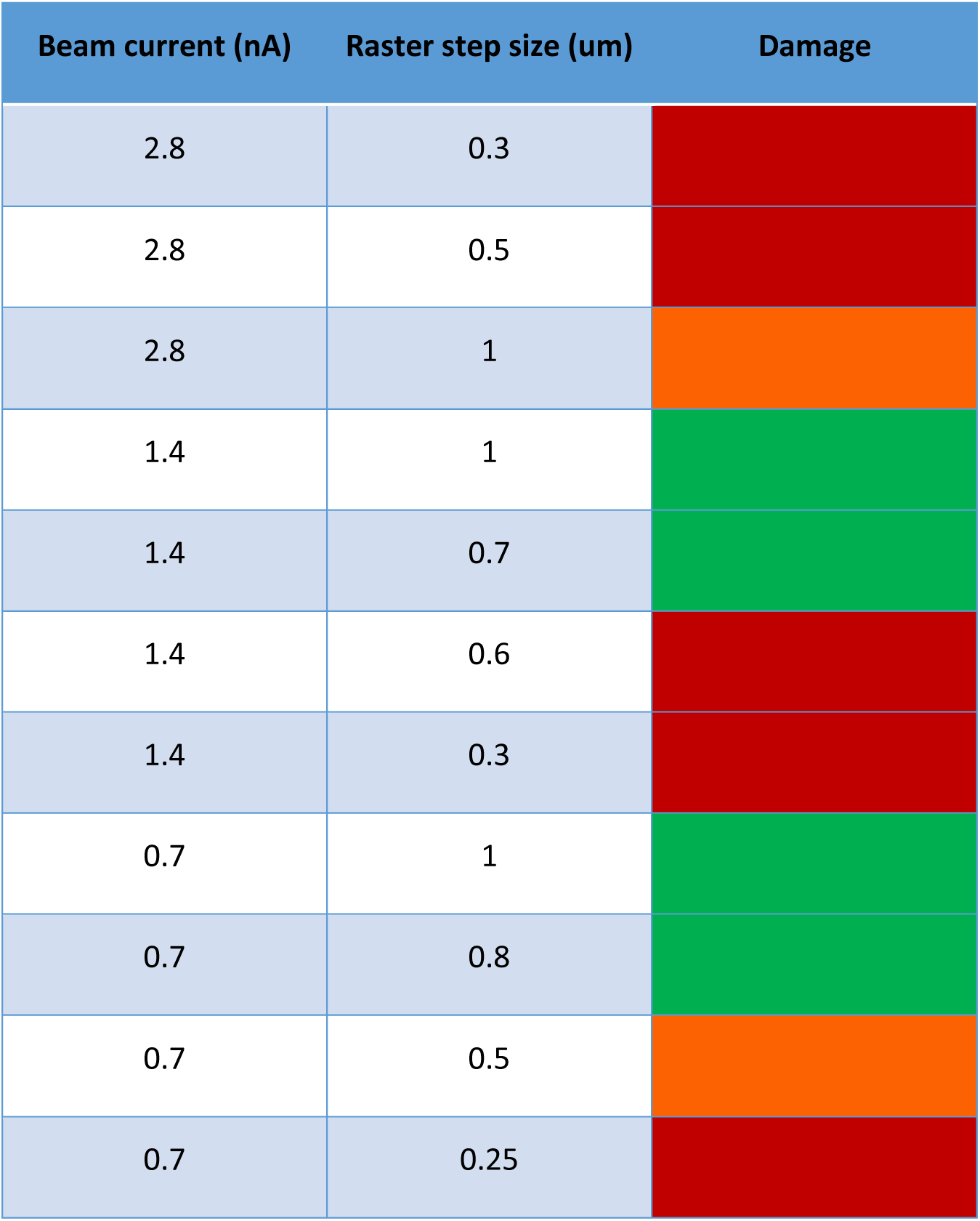
table corresponding to Figure 8, with red corresponding to measurable crystallization by EBSD, orange is damage only visible from SEM and not from EBSD, and green reflects no visible damage to the sample.

**Suppl. Figure 8:**
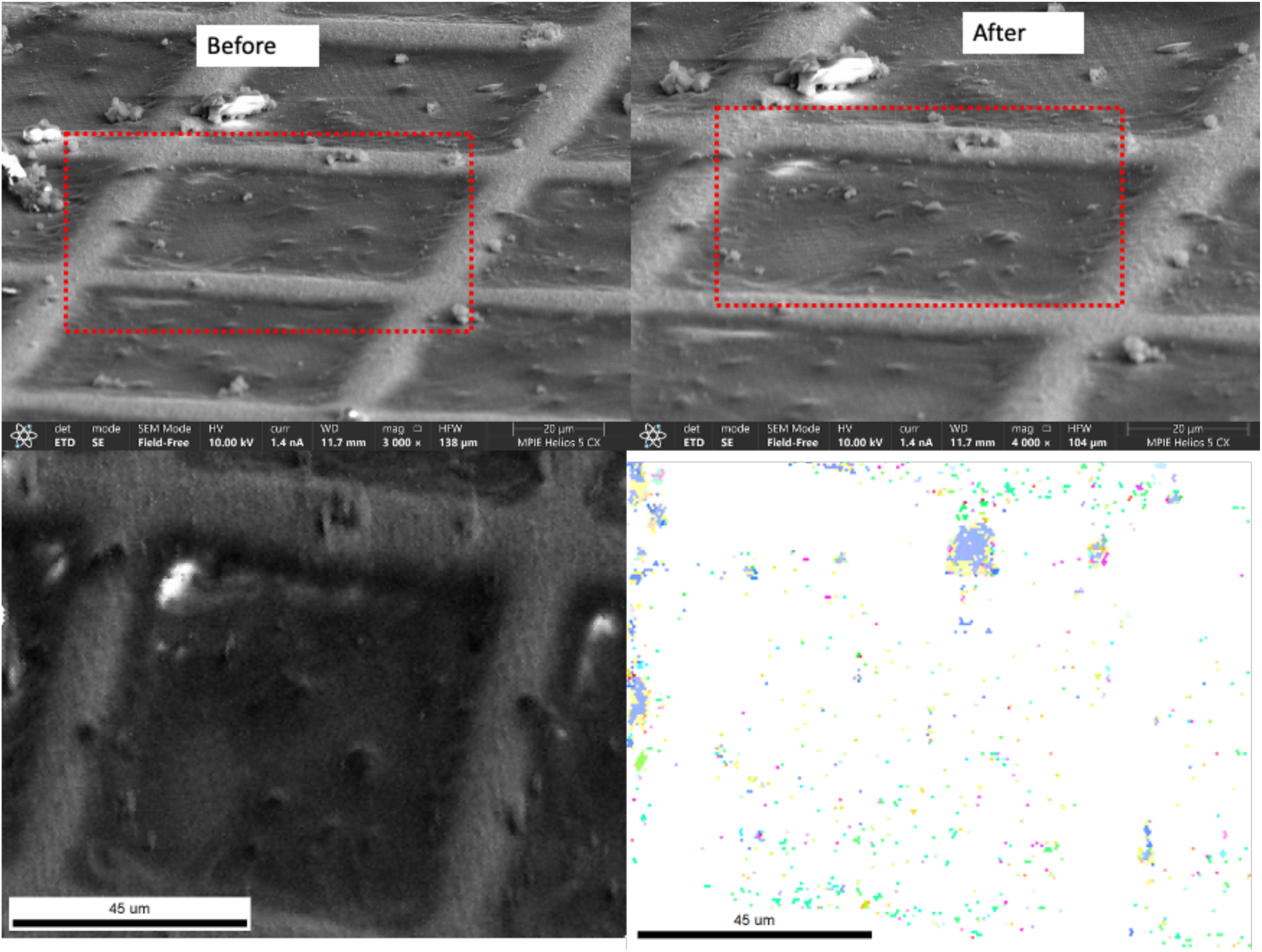
secondary electron micrograph of a cryo-TEM-grid carrying sea urchin sperm cells (a) before and (b) after an EBSD scan at 10kV, 1.4nA, 10kV, 2.8nA, and a 0.7 µm raster step size; corresponding (c) PRIAS image and (d) IPF from the EBSD scan indicating the presence of some ice crystals, including directly on the cells of interest.

**Suppl. Figure 9:**
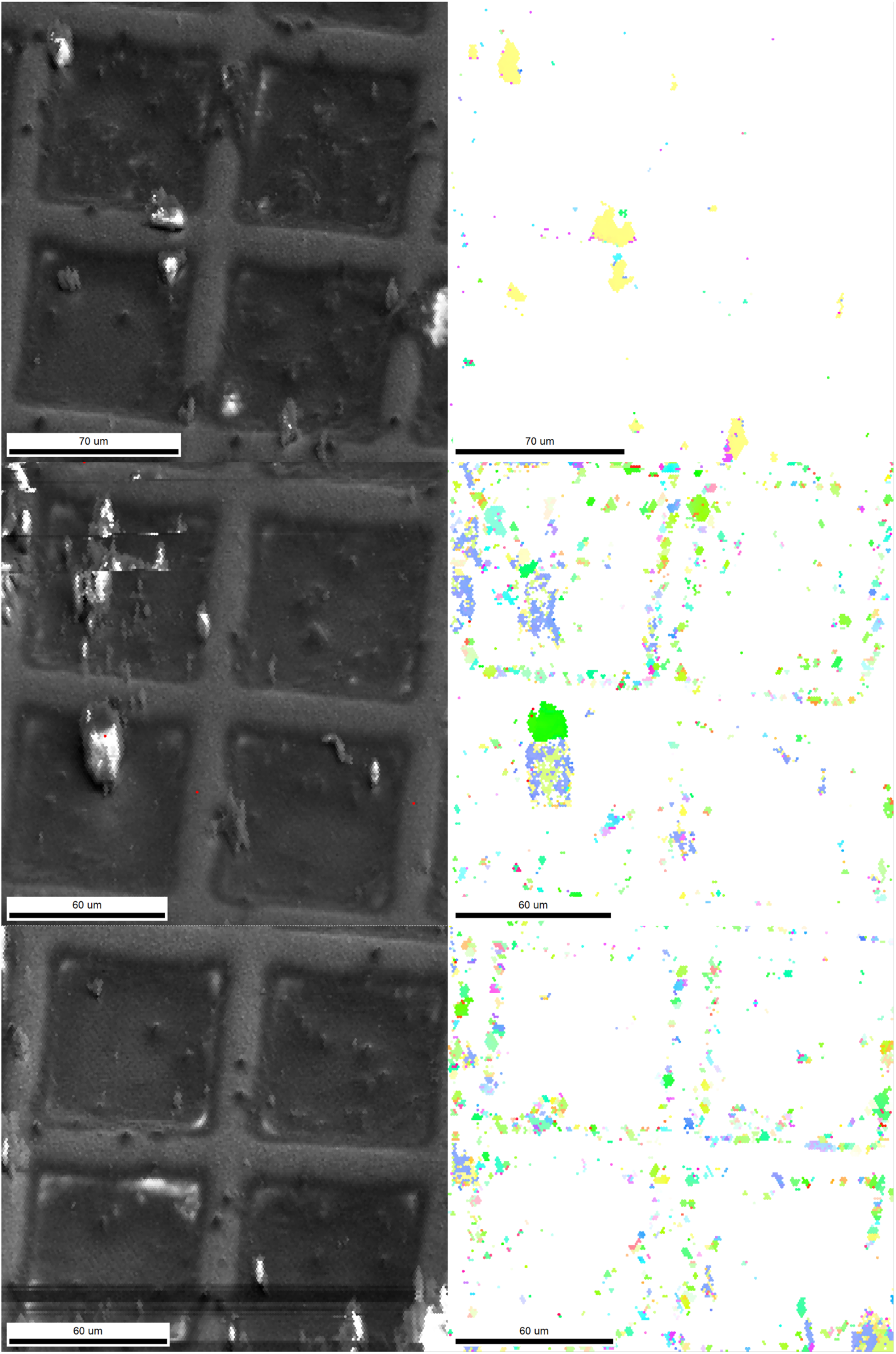
PRIAS and IPF for 3 different sets of 2×2 grid squares scans performed on the same grid with 10kV, 0.7 – 1pA, and 1um step size.

**Suppl. Figure 10:**
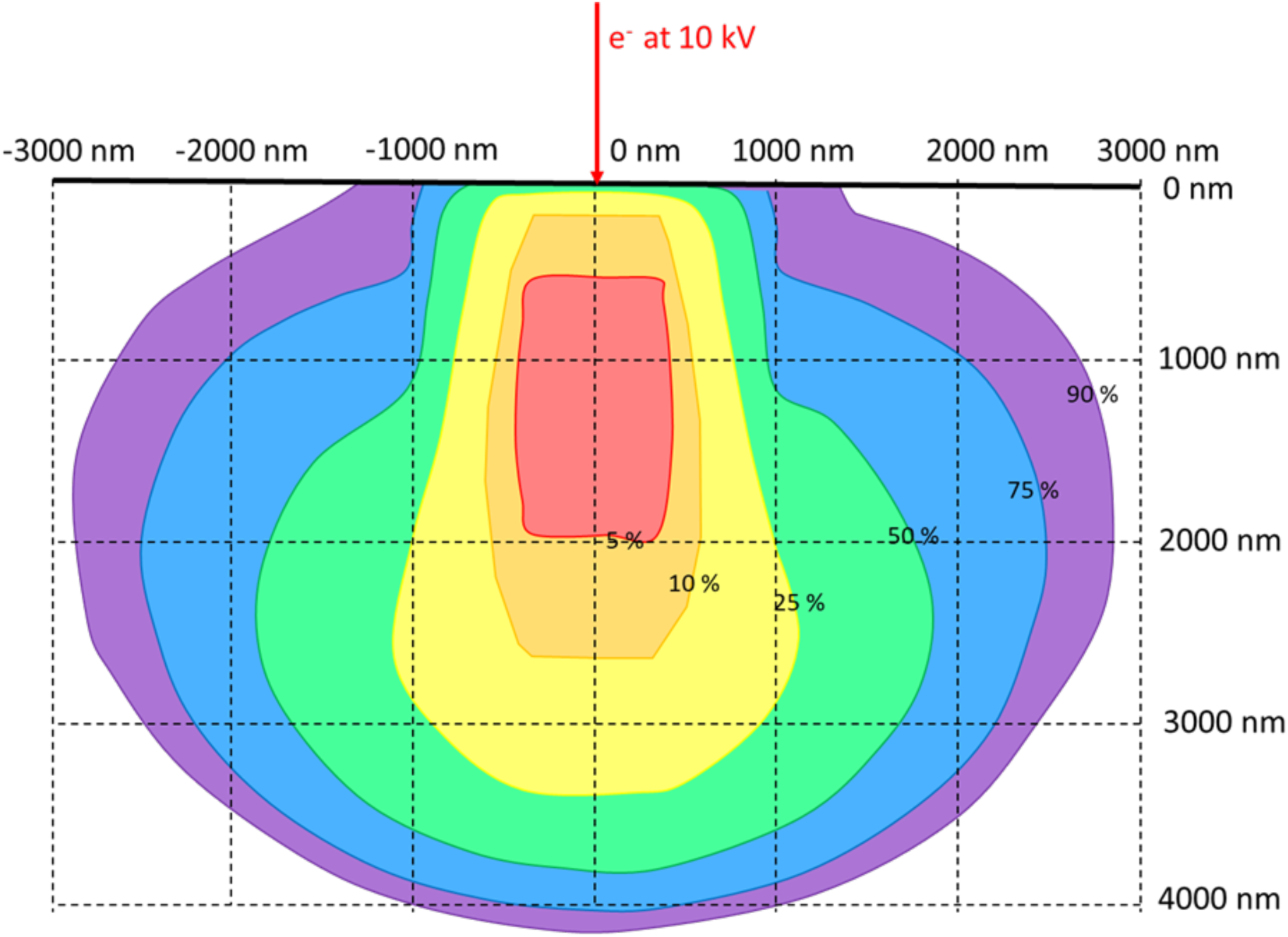
Casino Monte-Carlo trajectory simulation showing the total interaction volume for different energies with respect to the incident electron energy. The simulation is for 0° tilt (normal incidence) at 10 kV acceleration voltage. The red region corresponds to the volume where electrons have almost no energy loss, i.e. from where originate most backscattered electrons. The area where the most significant amount of energy (blue) is introduced into the material is near 5 µm in width.

**Suppl. Figure 11:**
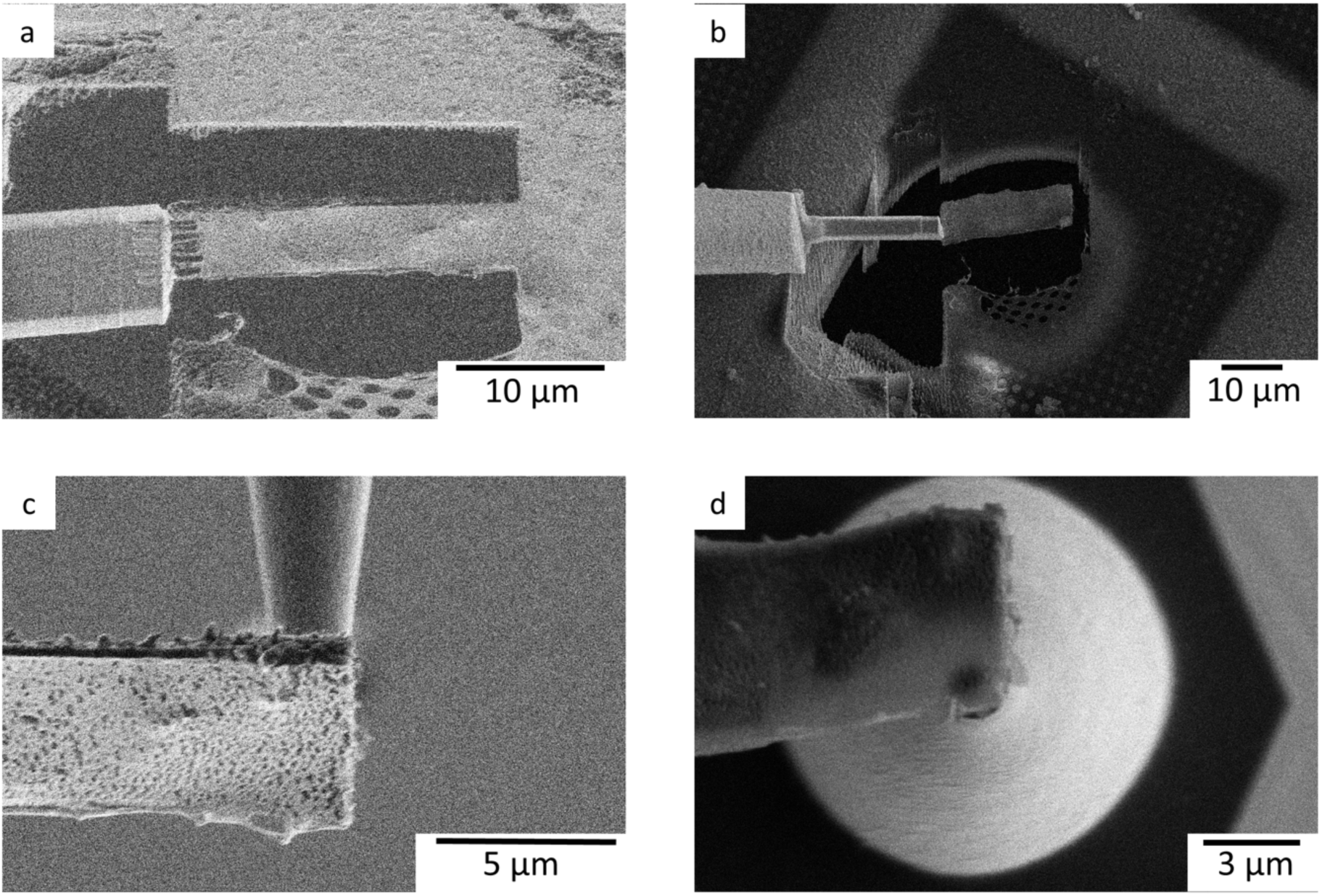
Proof-of-concept site specific lift-out from cryo-TEM grid to silicon coupon micropost. (a) FIB side view, redeposition welding of ice lamella from cryo-TEM grid (b) SEM top view, redeposition-welded lamella (c) FIB side view, lamella placed on Si microtip (d) SEM top view, lamella place on Si microtip.

